# Protein Surface Site Determines the Evolutionary Accessibility of Allosteric Regulation

**DOI:** 10.64898/2026.07.02.735819

**Authors:** Jerry C. Dinan, James W. McCormick, Rishi Soni, Samuel Thompson, Kimberly A. Reynolds

## Abstract

Domain recombination is a major source of new allosteric regulation in both evolved and engineered proteins. However, the sequence and structural features that govern where new allostery may emerge remain poorly understood. Here, we test the hypothesis that the evolutionary accessibility of allosteric regulation following domain insertion is constrained by local surface context, specifically association with pre-existing cooperative networks known as protein sectors. We began with two synthetic domain fusions wherein the *Avena sativa* light-oxygen-voltage (LOV2) domain was inserted into *Escherichia coli* dihydrofolate reductase (DHFR) at either a sector connected or non-sector connected surface. The insertion sites are only separated by five residues and both DHFR enzymes retain similar catalytic activity, yet the sector connected version exhibits a light-dependent allosteric phenotype, while the non-sector connected version does not. Using deep mutational scanning, we measured the effect of nearly all single point mutations on allostery in each chimera. The sector-connected DL121 was significantly more evolvable, possessing numerous allostery-tuning single mutants. In contrast, DL116 lacked statistically significant mutants that introduce allosteric regulation, suggesting the protein surface used by DL116 may be an evolutionary “dead end” for a regulatory phenotype. Surprisingly, DL116 did not show cooperative unfolding at temperatures up to 80 °C, suggesting that enhanced protein stability does not promote the evolvability of allosteric regulation as it does with other phenotypes. Together, our findings show that protein surface context influences the mutational pathways available for allosteric regulation, consistent with the view that sector-connected surface sites harbor a latent capacity for allostery while other locations are more evolutionarily inert.

## 1. Introduction

Domain recombination is a dominant source of novelty in protein evolution (Jin et al. 2009; Apic and Russell 2010; Peisajovich et al. 2010). Analyses of prokaryotic and eukaryotic genomes suggest that a majority of proteins are recombinants of multiple domains, accounting for about 65% of proteins in prokaryotes and 80% in eukaryotes (Apic et al. 2001). Domain fusion can generate novel function by altering localization, changing interaction partners, or conferring signaling phenotypes by establishing allosteric regulation between domains (Kuriyan and Eisenberg 2007; Jin et al. 2009; Peisajovich et al. 2010). Among the many forms of domain recombination, domain insertion appears particularly well suited to generate novel allosteric regulation. By embedding the inserted domain at a specific location within the host protein, domain insertion can direct conformational perturbations to a defined surface of the host domain, rather than relying on potentially transient interdomain contacts mediated by flexible linkers as is often the case in terminal fusions. Domain insertion has become a common strategy to engineer genetically encoded allosteric biosensors, including the proximity labeling tool LOV-Turbo (Lee et al. 2023), light or ligand-controlled kinases (Dagliyan et al. 2016; Dagliyan et al. 2017; Shaaya et al. 2020), phosphatases (Hongdusit et al. 2022), GTPases (Dagliyan et al. 2016), GEFs (Dagliyan et al. 2019), transferases (Reynolds et al. 2023), and transcription factors (Romano et al. 2021; Cao and Toettcher 2025). Inserting circularly permutated fluorescent proteins into sensor domains has also yielded a panoply of biosensors for sensing of Ca^2+^ (Nakai et al. 2001), glutamate (Marvin et al. 2013), GABA (Marvin et al. 2019), dopamine, acetylcholine (Jing et al. 2020), serotonin (Unger et al. 2020; Wan et al. 2021; Kubitschke et al. 2022), norepinephrine (Feng et al. 2019), neuropeptides (Wang et al. 2023), adenosine (Wu et al. 2023), ATP (Arai et al. 2018; Marvin et al. 2024), NADH (Hung et al. 2011; Zhao et al. 2015), NADPH (Tao et al. 2017), H_2_O_2_ (Belousov et al. 2006; Ermakova et al. 2014; Pak et al. 2020), pH, cAMP (Harada et al. 2017; Liu et al. 2022; Wang et al. 2022), membrane voltage (Chamberland et al. 2017; Lee and Bezanilla 2017; Xu et al. 2017), and many other signals (Greenwald et al. 2018; Kostyuk et al. 2019).

The typical domain insertion engineering workflow begins with generating an insertion variant library in which a sensory domain that detects light or ligand is inserted at various surface-exposed positions of a host protein, usually an enzyme or fluorescent protein (Stein 2017; Dagliyan et al. 2019) (Figure 1A). Insertion variants are then screened for regulatory phenotypes, in which host protein activity is modulated by the state of the sensory domain. Protein activity can be diminished or entirely lost following domain insertion. Even when function is preserved, initial regulatory phenotypes can be rare and show small dynamic ranges. Additionally, the direction of any emergent regulatory effect may not align with the specific goals of an engineering project. Typical engineering workflows thus follow an empirical, brute-force process of trial and optimization. Many fusions are constructed, and once an insertion site that produces an allosteric effect is identified, regulation is optimized through targeted mutagenesis, linker-length tuning, and/or directed evolution (Guntas and Ostermeier 2004; Nadler et al. 2016; Zhu et al. 2023). In contrast, insertion variants that fail to show an initial regulatory effect are usually discarded. Consequently, little is known about the comparative evolvability of allostery at distinct protein surfaces. These null sites may be fundamentally incapable of evolving allostery, or they may have access to monotonic single mutant paths towards allosteric regulation. Under selection for allostery, this can be conceptualized as a fitness landscape, where peak height corresponds to allosteric dynamic range, insertion site initializes a chimera’s location in sequence space, and the local topography of the fitness function determines the chimera’s evolutionary fate (Figure 1B) (Wright 1932; Maynard Smith 1970; Papkou et al. 2023).

**Figure 1:**
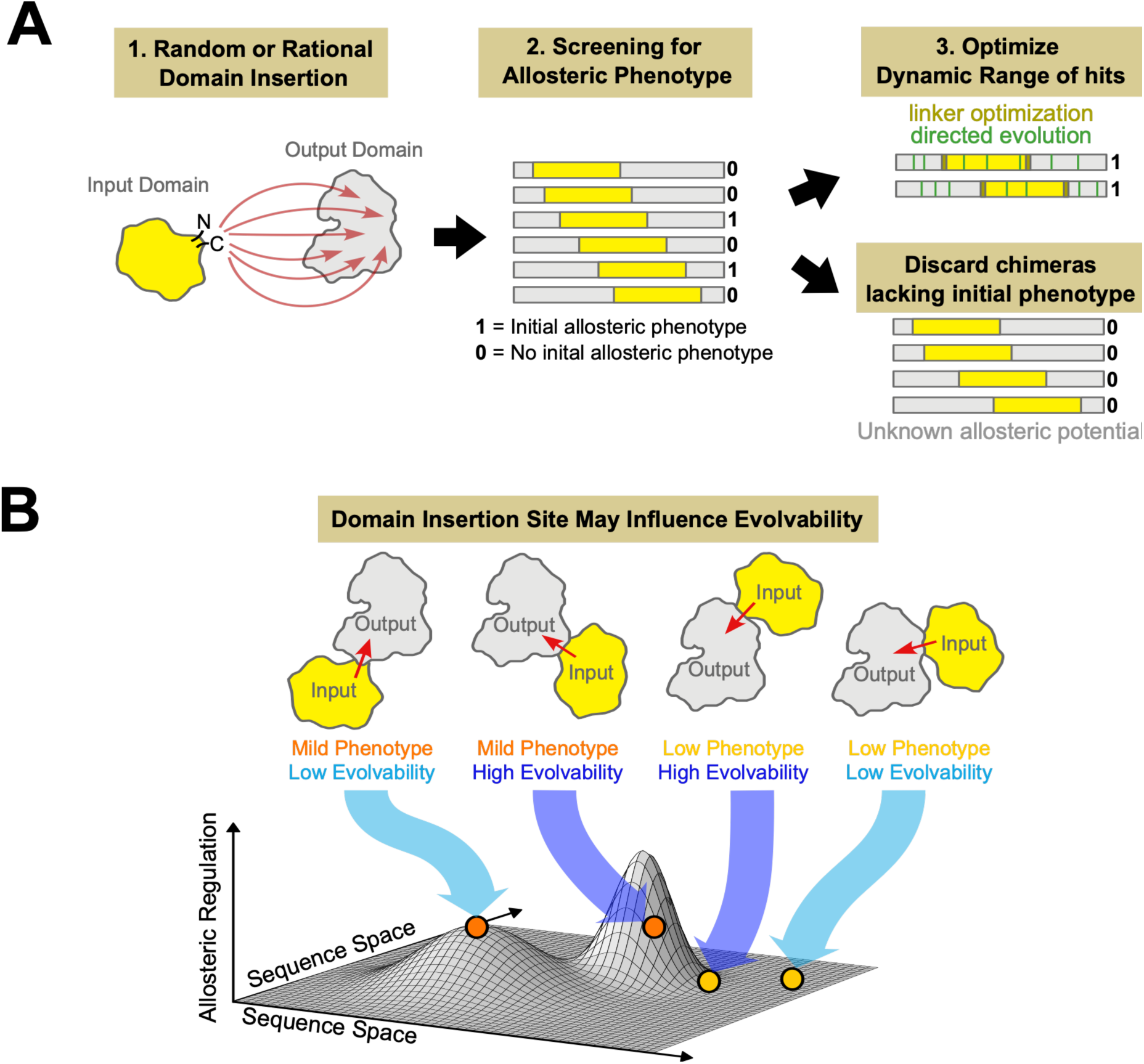
Domain insertion engineering allows for the interrogation of allosteric potential across protein surfaces. **(A)** The typical domain insertion engineering approach. An input domain is inserted into an output domain, or vice-versa. Screening is then performed to detect initial allosteric phenotypes in which the input domain regulates the output domain. Top allosteric hits are optimized, while chimeras lacking an initial phenotype are typically discarded. Therefore, little is known about the evolvability of allostery at initially inert sites. **(B)** Insertion of an “input” domain into an “output” domain can result in perturbations at different surfaces of the output domain. The insertion site influences initial allosteric phenotypes, but may also influence the evolvability of allosteric regulation, depending on where the resulting chimera exists on a fitness landscape.

This raises a central question: Is the evolvability of allostery uniformly distributed across a protein’s surface, or are some surface sites uniquely predisposed to evolve allosteric control? While some prior work has implied that all protein surface sites have allosteric potential (Gunasekaran et al. 2004), other studies have suggested that certain surface residues may in some way be primed with latent allosteric potential. Multiple lines of evidence suggest that energetic couplings within proteins are heterogeneous rather than uniform, leading to anisotropic mechanical properties and an uneven sensitivity to perturbation. For example, mutant cycle analysis in the Shaker potassium ion channel revealed energetic couplings across a sparse, defined network of residues (Sadovsky and Yifrach 2007). A consequence of these energetic coupling pathways is that energy transfer through proteins is highly dependent on the location of perturbation, as is evident in single-molecule pulling experiments in which protein unfolding is highly dependent on the location of applied force (Eyal and Bahar 2008). One framework for understanding energetic couplings and communication within proteins involves statistical analysis of intramolecular coevolution using methods such as statistical coupling analysis (SCA) (Lockless and Ranganathan 1999; Süel et al. 2003; Halabi et al. 2009; Rivoire et al. 2016). Here, the basic premise is that pairwise correlations in amino acid frequency across a diverse alignment indicate shared functional constraints. A key result of SCA is the identification of protein sectors, co-evolving residue groups that form contiguous networks of physical contacts connecting active or binding sites to distal surfaces (Lockless and Ranganathan 1999; Süel et al. 2003; Halabi et al. 2009; Smock et al. 2010). Protein sectors have been found to connect functional and allosteric sites in TonB-dependent transporters (Ferguson et al. 2007) and cryptochromes (Rosensweig et al. 2018). While not all distal sector-connected surfaces are used in nature as allosteric sites, these surfaces have been implicated as having latent allosteric potential. Experimental evidence from domain insertion scans, engineered phosphorylation in kinases, and small molecule drug binding suggests that perturbations at sector-connected surface sites are more likely to regulate protein function than perturbation at non-sector-connected sites (Lee et al. 2008; Reynolds et al. 2011; Novinec et al. 2014; Pincus et al. 2018). Together, these studies have inspired a model in which the evolution of new allosteric regulation is facilitated by pre-existing networks of energetically coupled residues as described by the sector. A logical but non-trivial consequence of this idea is that sector-connected domain insertions should likewise be readily tuned and optimized by virtue of connection at a cooperative network (Pincus et al. 2017). In turn, non-sector connected insertions would be refractory to further optimization, because establishing communication would require building cooperative interactions *de novo* across multiple amino acids.

Here, we test the hypothesis that allosteric regulation is more evolvable following domain insertion at sector-connected sites than at non-sector-connected sites. To do so, we compared mutational effects on allostery in two chimeras composed of the same sensory and host domains, differing by whether the sensory domain is inserted at a sector-or non-sector-connected site in the host domain. For the inserted sensory domain, we used the light-oxygen-voltage (LOV2) domain of *Avena sativa* Phototropin-1 kinase. The LOV2 domain is involved in activating signaling pathways involved in phototropism (Christie 2007; Hart and Gardner 2021). The flavin mononucleotide (FMN) chromophore of LOV2 absorbs a photon and reacts with the conserved C450 residue of LOV2, forming a covalent bond. In its native context, this triggers a conformational change in which the N-terminal A’α helix and the C-terminal Jα helix unfold. (Salomon et al. 2000; Crosson and Moffat 2002; Halavaty and Moffat 2007; Iuliano et al. 2020) For the host domain, we used *Escherichia coli* dihydrofolate reductase (DHFR). DHFR is an essential metabolic enzyme in folate metabolism that reduces dihydrofolate (DHF) to tetrahydrofolate (THF) with the concomitant oxidation of NADPH to NADP^+^. This activity is essential for the maintenance of folate pools and the biosynthesis of purines and some amino acids (Schober et al. 2019). The strong dependence of *E. coli* growth on DHFR activity allows for mutational effects in DHFR-LOV2 chimeras to be interrogated with high throughput growth measurements (Reynolds et al. 2011; McCormick et al. 2021). Contrasting the growth rate effects of mutations in the light and dark allows mapping of the local fitness landscapes of allosteric regulation for sector-and non-sector-connected chimeras. Leveraging this property, we measured the mutational effect of nearly all possible single mutants of DHFR on allostery for a representative non-sector connected and sector connected DHFR-LOV2 fusion. The sector-connected fusion can drive a 35% increase in growth rate in the light versus the dark, and we observed statistically significant allosteric tuning for 52 mutations. In contrast, the non-sector-connected DHFR-LOV2 chimera lacks an initial regulatory phenotype both *in vitro* and *in vivo* and lacks any single mutational path that significantly introduces allosteric regulation, despite having a similar baseline biochemical activity compared to the sector-connected chimera and having LOV2 inserted only five residues away. Biophysical measurements and structural modeling suggest that the non-sector connected domain insertion forms a stabilizing interface between the inserted and host domains, which may constrain the conformational changes required for allosteric coupling. Together, these results demonstrate that surface sites differ in their capacity to evolve allosteric regulation even if domain insertion is equally well-tolerated in terms of activity. Our findings are consistent with a model in which the evolution of allostery is facilitated by pre-existing pathways of cooperative residues within proteins described by coevolutionary sectors.

## 2. Results

### 2.1 Two structurally proximal DHFR/LOV2 fusions retain domain function but diverge in allosteric regulation

To establish a model system for comparing the evolvability of allostery across protein surfaces, we characterized two fusions in which LOV2 was inserted into the enzyme DHFR. Both insertions occurred in the βF-βG loop of DHFR, a loop that influences DHFR catalysis through dynamic interactions with the catalytic Met20 loop. One insertion occurred between DHFR positions 115 and 116 which are not part of the sector, resulting in a chimera termed DL116. The other insertion occurred between residue 120 and the sector residue 121, resulting in a chimera termed DL121. Prior *in vivo* functional assays indicated that DL116 is not allosterically regulated, while DL121 was previously shown to have a modest allosteric effect both i*n vivo* and *in vitro* (Lee et al. 2008; Reynolds et al. 2011). To more carefully assess sector contacts between the DHFR sector and the inserted LOV2 domain, we generated AlphaFold2 models of DL116 and DL121. DL116 was predicted to lack direct contacts between the DHFR sector and the terminal helices of LOV2 (Figure 2A), while DL121 exhibited physical contact between the DHFR sector and the Jα helix of LOV2 (Figure 2B). The DL116 chimera was also predicted to form an extensive interface with 1,889.89 Å^2^ of buried surface area between the LOV2 domain and the beta sheet face the major subdomain of DHFR, while the DL121 model indicated a smaller interdomain interface with 1,095.23 Å^2^ of buried surface area. The interdomain interfaces of DL116 and DL121 involve 224 and 105 atoms of DHFR, respectively, with only an 11-atom overlap. These structural models suggest that LOV2 switching in DL116 and DL121 would perturb unique surfaces of DHFR, despite LOV2 being inserted only 5 residues apart.

**Figure 2:**
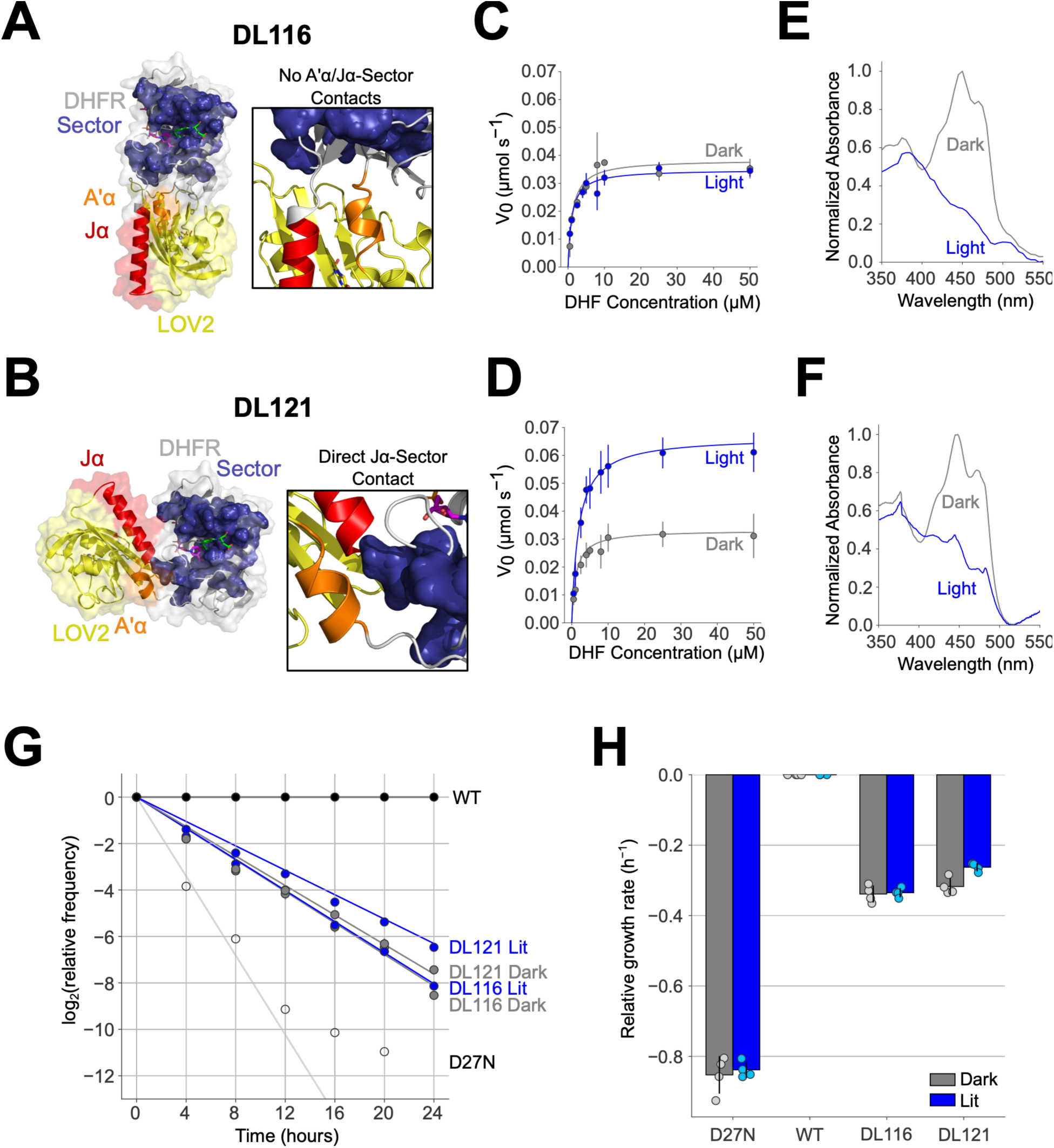
Engineered DHFR-LOV2 chimeras DL116 and DL121 represent functional nascent domain fusions. (A-B) AlphaFold2 models of (A) DL116, a chimera in which LOV2 is inserted between residues 115 and 116 of DHFR, and (B) DL121, a chimera in which LOV2 is inserted between residues 120 and 121 of DHFR. The backbone of DHFR is shown in grey cartoon, and the LOV2 domain in yellow cartoon. The Jα and A’α helices of LOV2 are highlighted in red and orange, respectively. Folate, NADPH, and FMN are shown as green, pink, and yellow sticks, respectively. (A) The DL116 chimera lacks a direct contact between the DHFR sector, shown as blue spheres, and the Jα and A’α helices. (B) The DL121 chimera has a direct contact between the end of the Jα helix of LOV2 and the DHFR sector. **(C-D)** Steady state kinetics for the two chimeras. Circles indicate the mean of triplicate measurements, with error bars showing the standard deviation. The blue and gray lines represent the best fit of the steady-state Michaelis Menten equation. (C) DL116 shows no significant difference in DHFR activity in the light (blue) and dark (gray). (D) DL121 shows a two-fold increase in DHFR activity following light exposure. Circles show the mean of triplicate measurements, and the error bars show the standard deviation across three replicates. **(E-F)** Absorbance spectra of (E) DL116 and (F) DL121 immediately following light exposure (blue) and after relaxation into the dark state (gray). The light-driven decrease in absorbance at 447 nm indicates that the LOV2 domain is functional in both chimeras. Absorbance spectra were min-max normalized within each construct to compare spectral state between lit and dark states. **(G)** Growth rates relative to WT DHFR in the light and dark were calculated using weighted least squares (WLS) linear regression of the log relative mutant frequency vs. time, weighted by sequencing counts. Datapoints are the mean over quadruplicate biological replicates. The error bars show standard deviations, which are in most cases obscured by the datapoints. Analogous data for the inactive DHFR-D27N mutant is shown for comparison. **(H)** Barplot of average relative growth rates of DHFR-D27N, DL116, and DL121 in the light and dark determined by WLS linear regression. Error bars show the standard deviation of the growth rate. Individual measurements are shown as circles. DL116 has a relative growth rate of −0.34 h^−1^ in the dark and −0.34 h^−1^ in the light, with a non-significant allosteric effect (p = 0.797). DL121 has a relative growth rate of −0.32 h^−1^ in the dark and −0.26 h^−1^ in the light, with a significant allosteric effect of 0.0547 h^−1^ (p = 0.0116).

To further compare the initial levels of allosteric regulation of DHFR by LOV2 in both chimeras, we measured steady state DHFR kinetics following white light exposure, or in the dark (Table S1). The non-sector-connected DL116 construct did not exhibit significant light-dependent kinetics *in vitro*, with a dark k_cat_ of 0.38 ± 0.03 s^−1^, a lit k_cat_ of 0.349 ± 0.001 s^−1^ (p = 0.26), a dark K_M_ of 1.5 ± 0.5 μM, and a lit K_M_ of 1.2 ± 0.1 μM (p = 0.07) (Figure 2C). In contrast, the sector-connected DL121 construct displayed a 2-fold increase in k_cat_ following light activation, with a dark k_cat_ of 0.33 ± 0.07 s^−1^, a lit k_cat_ of 0.67 ± 0.07 s^−1^ (p = 0.007), a dark K_M_ of 1.5 ± 0.3 μM, and a lit K_M_ of 2.1 ± 0.3 μM (p = 0.05) (Figure 2D). DL121 thus exhibits V-type allostery (Carlson and Fenton 2016). Spectroscopic characterization of LOV2 activation found that both chimeras exhibited the characteristic loss of absorbance at 447 nm following light exposure (Figure 2E-F), indicating light-induced cysteinyl-flavin adduct formation. The absence of allosteric regulation in DL116 can therefore not be attributed to impaired photosensitivity of its LOV2 domain. We used an NGS-based growth assay to measure growth rates of each chimera relative to WT DHFR (McCormick et al. 2021) (Figure 2G). DL116 exhibited a non-significant change in growth rate in response to light (p = 0.797). Conversely, light exposure increased the relative growth rate of the DL121 strain from −0.32 h^−1^ to −0.26 h^−1^ (p = 0.0116). Both chimeras exhibited similar growth rates in the dark, with growth rate phenotypes between wild-type DHFR and the catalytically inactive DHFR mutant D27N (Figure 2H). These findings confirm that: (1) the two chimeras retain comparable levels of baseline DHFR activity both *in vitro* and *in vivo* and (2) the LOV2 insertion site influences the initial regulatory phenotype (Reynolds et al. 2011). Additionally, recent NMR and biochemical characterization of DL121 from our lab suggests that the order to disorder transition in LOV2 drives localized conformational changes in the DHFR active site that are linked to the transition state free energy of catalysis (McCormick et al. 2024). Together these data establish DL121 as a nascent allosteric switch, while DL116 lacks productive communication between domains.

### 2.2 DL121 exhibits greater mutational robustness than DL116

We next compared the evolvability of allosteric regulation in the two fusions using deep mutational scanning (Fowler and Fields 2014). We had previously constructed a saturation mutagenesis library across all positions of the DHFR domain of DL121 and measured the impact of nearly all single mutants on growth rate in the light and the dark (McCormick et al. 2021). For this work, we constructed an analogous saturation mutagenesis library of the DHFR domain of DL116. The DL116 library was transformed into an *E. coli* strain lacking endogenous genomic DHFR (ER2566 Δ*folA* Δ*thyA*), and the resulting cells were selected in M9 minimal media under lit and dark conditions. As in our previous work, we used next generation sequencing to monitor the frequency of each variant over time. To ensure an apples-to-apples comparison of our DL116 deep mutational scan to the previous scan of DL121, we performed downsampling of the sequencing data to ensure matched read depth across the two experiments (Supplementary Figure S1). After downsampling, the initial sequencing timepoints contained 3,091 of the 3,180 mutants in the DL116 library (97.20% complete) and 3,047 of 3,180 mutants in the DL121 library (95.82% complete) (Supplementary Figure S2A-B). The sequencing counts per mutant had roughly log-normal distributions (Supplementary Figure 2C-D). We then used weighted linear regression to fit relative growth rates to the variant frequencies over time (Rubin et al. 2017) (see details in methods) (Figure S3, Table S2). The use of downsampled reads and weighted linear regression led to modestly different relative growth rates in comparison to our original analysis of the DL121 dataset (Supplementary Figure S4).

The distribution of relative fitness effects was qualitatively similar across the two chimeras, and between lit and dark conditions. The mean growth rate of DL116 mutants was −0.07 h^−1^ in the dark and −0.06 h^−1^ in the light, while the average for DL121 was −0.1 h^−1^ in both dark and light. Both distributions indicate that the majority of mutations are deleterious relative to the starting chimera with a handful of mutations that are beneficial. This observation contrasts to many native proteins, where the vast majority of mutations are often near neutral (McLaughlin Jr et al. 2012; Thompson et al. 2020; Nguyen et al. 2024). Because DL116 and DL121 already exhibit reduced activity relative to wild-type DHFR, they are expected to reside near the steep portion of the DHFR activity–fitness relationship. Consequently, small decreases in activity can cause substantial reductions in growth rate, making deleterious mutations common in our assay. This same sensitivity also enables the detection of modest light-dependent changes in activity.

To determine the relative mutational robustness of the two chimeras, we defined a viability cutoff as the mean growth rate of nonsense mutants in the first 80 residues of DHFR. Mutants with a growth rate below this cutoff likely reflect noisy measurements of variants with low or no DHFR activity and are thus excluded from subsequent analysis. For mutants above this viability cutoff, we found a high level of inter-replicate reproducibility (mean R^2^ of 0.90 in the dark and 0.92 in the light for DL116 and mean R^2^ of 0.96 in the dark and in the light for DL121) (Supplementary Figure S5A-D). In the dark, 59% of DL116 mutants and 74% of DL121 mutants were viable. And in the light, 59% of DL116 mutants and 73% of DL121 mutants were viable (Figure 3A-B). Thus, the DL121 chimera was more mutationally robust. In DL116, non-viable mutants were enriched at 34 positions in the dark and 28 in the light. In DL121, non-viable mutants were enriched at 12 positions in the dark and 15 in the light (Figure 3C-D). This means that fewer positions in DL121 were especially sensitive to mutation, consistent with DL121 being more mutationally robust overall. Non-viable mutations in DL121 were primarily located around the active site, while in DL116 they also occurred in the β-sheets of the major subdomain. Some of these mutationally sensitive positions in DL116 are proximal to the proposed interface with LOV2. Moreover, non-viable mutations were significantly enriched in the DHFR sector in both chimeras in both the light and dark (Supplementary Figure S6), consistent with the view that residues in the sector are important for DHFR activity.

**Figure 3:**
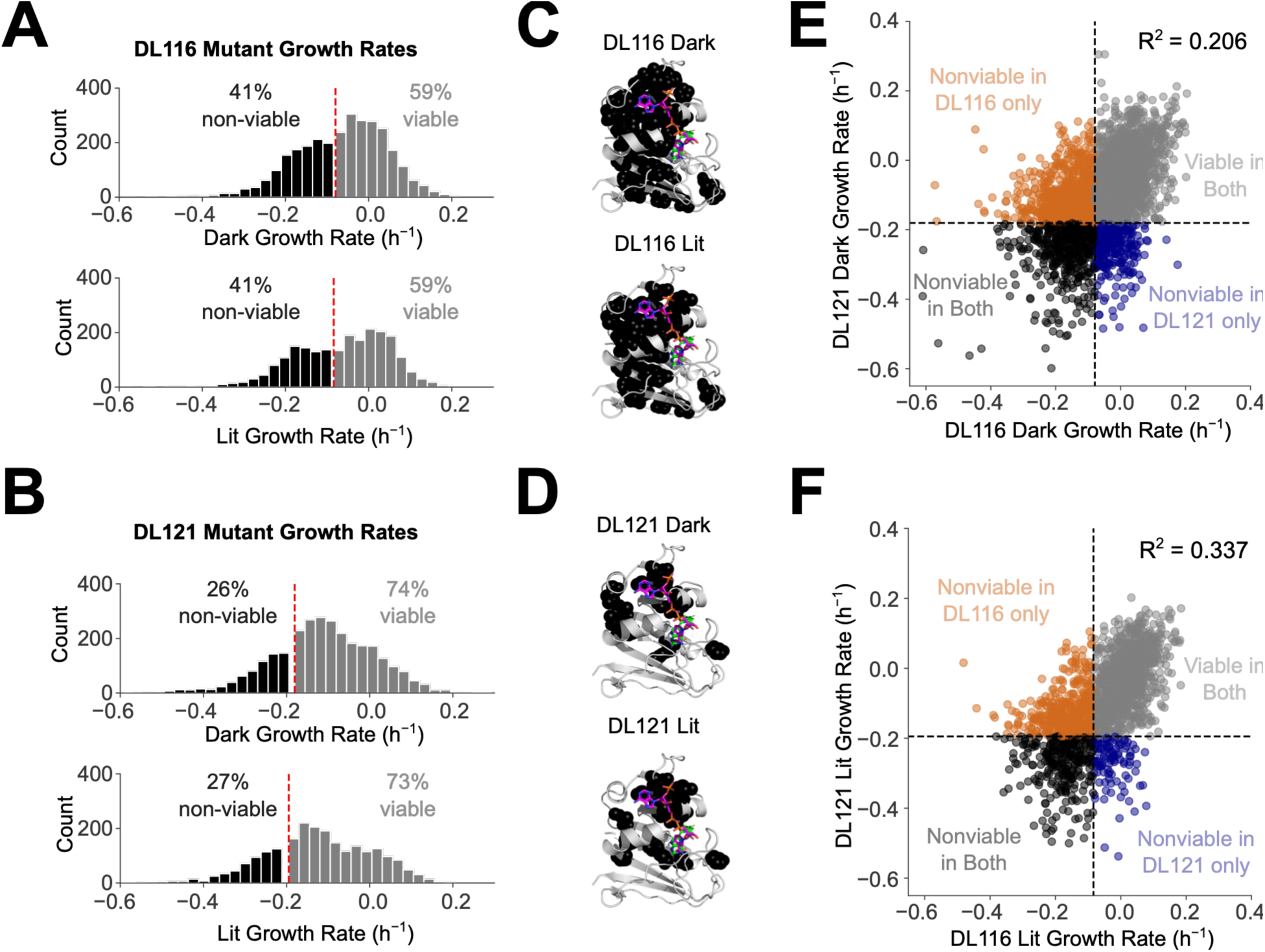
DL121 is more mutationally robust than DL116. (A-B) Distributions of mutational effects on growth rate in DL116 (A) and DL121 (B) in the dark (upper) and light (lower). The red dashed lines show viability cutoffs equal to the mean growth rates of nonsense mutations at positions 1-80 of DHFR. **(C-D)** Positions enriched for non-viable mutations shown as black spheres on a crystal structure of DHFR (PDB: 1RX2). NADPH is shown in pink and folate in green. **(E-F)** Correlations of mutational effects in the DL116 and DL121 chimeras in the dark (E) and light (F). Each point represents the mean of triplicate measurements. Viability cutoffs are shown as black dashed lines.

We observed a modest correlation between the relative growth rates of DL116 and DL121 mutants (R^2^ of 0.206 in the dark and 0.337 in the light) (Figure 3E-F). The correlated mutational effects imply that the chimeras share some similarities energetically, but the insertion site likely has an epistatic influence on mutational effects. In the dark, 49.27% of mutations were viable in both chimeras, 10.06% were viable in DL116 only, and 25.05% were viable in DL121 only. In the light, 54.39% of mutations were viable in both chimeras, 6.385% of mutations were viable in DL116 only, and 23.38% of mutations were viable in DL121 only.

### 2.3 Allosteric regulation is more evolvable in DL121 than in DL116

Next, we determined the effect of each mutation on allosteric regulation in DL116 and DL121. We subtracted each mutant’s relative growth rate in the dark from its relative growth rate in the light to calculate allostery tuning effects (ΔA) (Supplementary Figure S7A-C). In NGS-based measurements of relative growth rate, each mutation is assigned a growth rate relative to a particular reference sequence. Consequently, the ΔA allostery-tuning scores are by definition normalized such that each unmutated chimera has a ΔA score of zero. Because DL116 and DL121 have different initial levels of allostery, we implemented a “baseline correction” to account for this initial, or baseline, allostery (A_0_). To convert the relative allostery tuning effects to absolute allosteric dynamic ranges (ADR), we added the baseline allosteric effect sizes to each score (Supplementary Figure S7A, S7D-E). Because unmutated DL116 does not exhibit detectable allostery, mutants introducing allostery were classified as either positive or negative. Positive mutants introduced a phenotype in which growth increased in the light, whereas negative mutants introduced a phenotype in which growth decreased in the light. In contrast, DL121 already possesses positive allostery, allowing mutants to be classified as enhancing or disrupting based on their effects on the magnitude and direction of light-dependent regulation.

The majority of viable mutants had relatively small effects on allostery, with unimodal distributions centered around the original phenotypes of each chimera (Supplementary Figure S7F). In DL116, the distribution of allosteric dynamic range (ADR) appeared slightly skewed in the positive direction with 59.7% positive allosteric mutants, and 40.3% negative. In DL121, 99.2% of mutants appeared to retain positive allostery, with 63.4% showing reduced ADR and 35.8% showing enhanced ADR. The remaining 0.8% of viable DL121 mutants apparently reversed the direction of regulation to negative allostery. We constructed volcano plots for each chimera to assess the statistical significance of these allostery-tuning effects. In DL116, 47 of 1,253 viable mutants (∼4%) displayed light-dependent growth with p < 0.05 (Figure 4A), whereas in DL121, 148 of 1,632 viable mutants (∼9%) displayed light-dependent growth with p < 0.05 (Figure 4B). We used the sequential goodness of fit (SGoF) algorithm to correct for multiple hypothesis tests (Carvajal-Rodríguez et al. 2009). After multi-test correction, none of the DL116 mutants exhibited statistically significant allostery, whereas 52 DL121 mutants (∼3%) remained significant. Because differences in noise between datasets could influence statistical power (Supplementary Figure S5), we performed a label-permutation test to compare the overall enrichment of allostery-tuning mutants in each chimera (Good 1994). P-values for mutants across both datasets were pooled and randomly reassigned to DL116 or DL121 while preserving dataset size, and the number of mutants with p < 0.05 was recorded for each permutation. The observed number of mutants with p < 0.05 in DL121 was significantly enriched relative to the permuted null distribution (p = 4 x 10^−9^), whereas DL116 showed a correspondingly significant depletion (Figure 4C-D). Although differences in noise may influence statistical power, the unequal distribution of low p-values between DL116 and DL121 is highly unlikely to have arisen by chance. Taken together, these analyses support the conclusion that mutations are more likely to tune allosteric regulation in the sector-connected DL121 chimera than in the non-sector-connected DL116 chimera.

**Figure 4:**
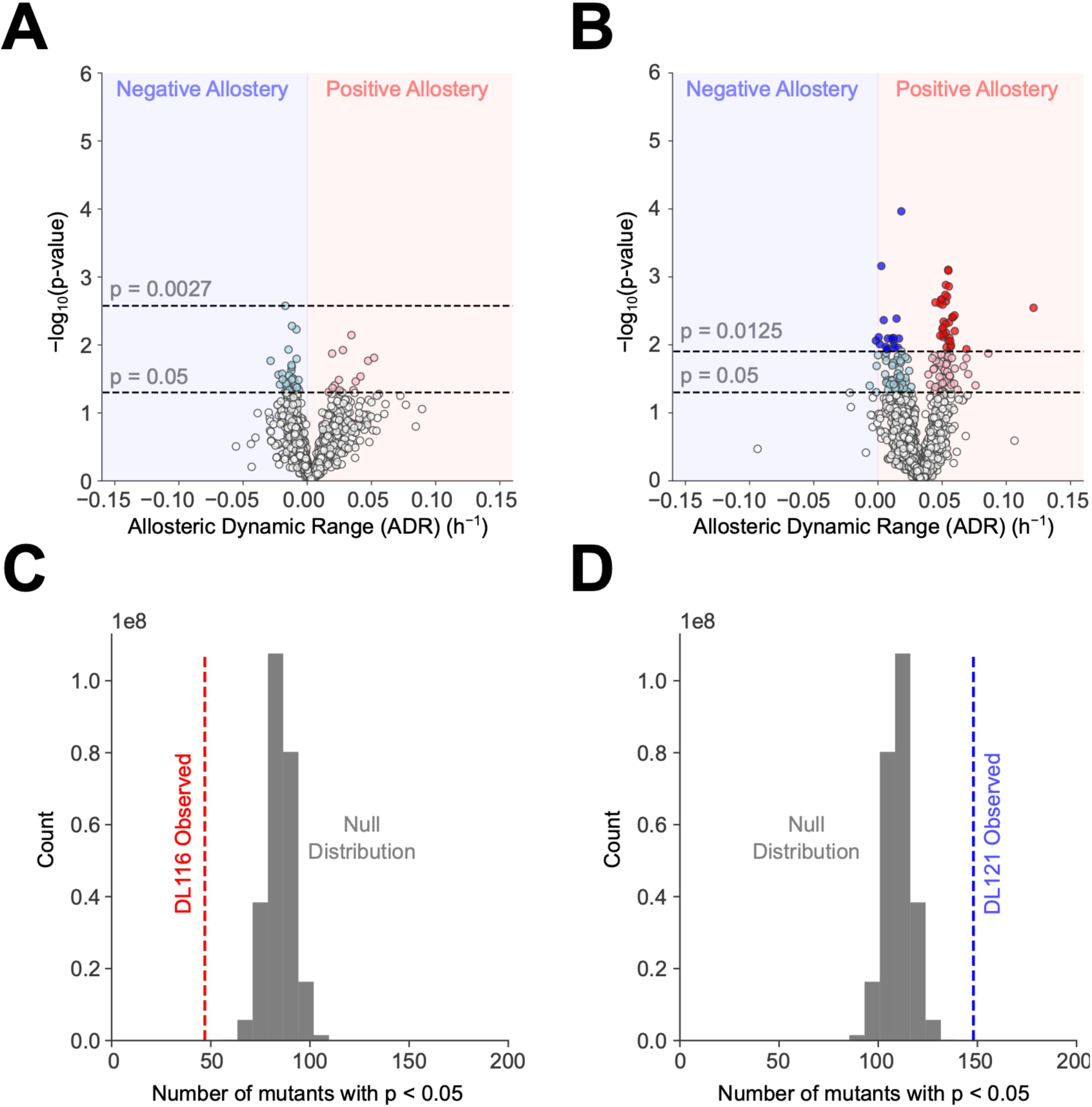
Allosteric regulation is more evolvable in DL121 than in DL116. **(A)** Volcano plot of allosteric dynamic range for DL116 mutants. Nominally-significant mutants (p < 0.05) that introduce either negative or positive allosteric regulation are shown in light blue and pink, respectively. The standard p-value cutoff (p = 0.05) and the SGoF-adjusted cutoff (p = 0.0027) are shown as black dashed lines). **(B)** Volcano plot of allosteric dynamic range for DL121 mutants. Nominally-significant mutants that disrupt or enhance allosteric regulation are shown in light blue or pink, respectively. Mutants that pass SGoF multiple hypothesis correction and significantly disrupt or enhance allosteric regulation are shown in blue or red, respectively. The standard p-value cutoff (p = 0.05) and the SGoF-adjusted cutoff (p = 0.0125) are shown as black dashed lines). **(C-D)** Label-permutation enrichment tests of allostery-tuning p-values associated with DL116 (C) or DL121 (D) mutants. Null distributions of the number of mutants with p < 0.05 following 250,000,000 permutations are shown as grey histograms. The observed number of mutants with p < 0.05 are shown as dashed lines.

### 2.4 Domain insertion site influences the structural distribution of allostery-tuning mutations

To compare the structural distributions of allostery-tuning mutations in DL116 and DL121, we mapped mutants exhibiting nominally significant allostery (p < 0.05) and multiple-hypothesis-corrected allostery (p < 0.0125 for DL121) onto AlphaFold models of each chimera. In DL116, positive allosteric mutations with p < 0.05 appeared clustered near the NADPH-binding cleft in the adenosine-binding subdomain and around the DHFR–LOV2 domain interface in the major subdomain (Figure 5A). Some of the negative allosteric mutants of DL116 with p < 0.05 formed a cluster or “bridge” connecting the Jα helix to the DHFR sector (Figure 5B). These bridge mutants may indicate routes through which conformational changes in LOV2 can be coupled to sector residues in DHFR. These structural distributions suggest that cofactor interactions or alteration of interdomain contacts may allow for the emergence of light-dependent regulation in DL116. For DL121, allostery-tuning mutants with p < 0.05 were distributed across the surface of DHFR, as previously reported in McCormick et al. 2021 (Figure 5C-D) (McCormick et al. 2021). Allostery-disrupting mutants of DL121 that passed SGoF multi-test correction appeared concentrated near the LOV2 insertion site (Figure 5D) (McCormick et al. 2021). The contrasting structural distributions in DL116 and DL121 suggest that insertion site not only affects the magnitude of initial allosteric phenotypes, but also shapes the mutational paths available for subsequent optimization of allostery. The large distribution of allostery-tuning mutants across the DL121 structure may mean that DL121 has a high degree of global cooperativity, perhaps due to the allosteric perturbation being directed to the sector. This contrasts with DL116, in which we see weak allostery-tuning mutants concentrated in specific regions of the protein, possibly consistent with the protein needing to establish new couplings between LOV2 and the DHFR sector before making true gains in allostery.

**Figure 5:**
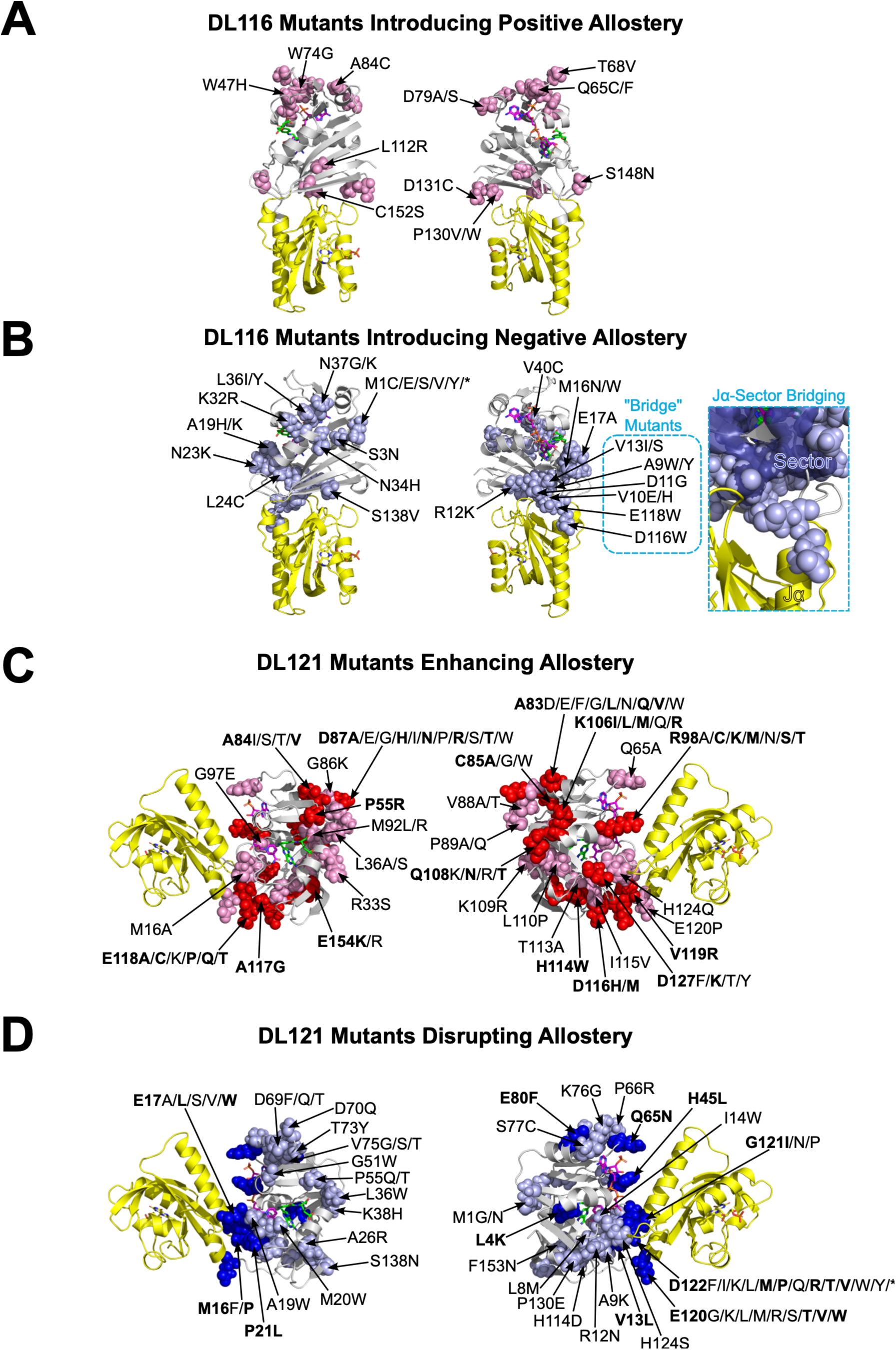
Domain insertion site influences the structural distribution of allostery-tuning mutations. In each panel, the LOV2 domain is shown in yellow cartoon and the DHFR domain is shown in grey cartoon. NADPH, folate, and FMN are shown as pink, green, and yellow sticks, respectively. **(A)** Two views of DL116; positions harboring mutations that introduce nominally-significant positive allosteric regulation are shown in pink (p < 0.05). **(B)** Two views of DL116; positions harboring mutations that introduce nominally-significant negative allosteric regulation are shown in light blue (p < 0.05). A zoomed-in view of the allostery-introducing mutants that “bridge” the end of the Jα helix of LOV2 and the DHFR sector is shown to the right. **(C)** Two views of DL121, with positions harboring mutations that enhance existing allosteric regulation shown in either pink (nominally significant, p < 0.05) or red (significant, passing SGoF-adjusted cutoff of p = 0.0125). Labels for mutants passing multiple hypothesis correction are bold. Several positions contain multiple significant allostery-enhancing mutants. **(D)** Two views of DL121, with positions harboring mutations that enhance existing allosteric regulation shown in either light blue (nominally significant, p < 0.05) or blue (significant, passing SGoF-adjusted cutoff of p = 0.0125). Labels for mutants passing multiple hypothesis correction are bold.

### 2.5 DL116 is a thermostable enzyme

The stability of enzymes has been proposed to influence their evolvability. Enhanced stability can buffer mutational variation that leads to improvements in activity of function at the expense of stability (Bloom et al. 2005; Bloom et al. 2006). To determine the relative stabilities of DL116 and DL121, we performed thermal melts while monitoring unfolding with circular dichroism spectroscopy. Prior to melting, we collected far-UV CD spectra at 20 °C. These spectra displayed negative circular dichroism near 220-225 nm, consistent with folded proteins (Figure 6A). DL121 had a slightly more negative mean residual ellipticity near 220 nm compared to DL116, possibly indicating the presence of more secondary structural content in DL121. DL121 appears to unfold and then aggregate when heated from 20 °C to 80 °C (Figure 6B). This is consistent with our previous work performing CD melts on DL121 (McCormick et al. 2024). In contrast, DL116 exhibited little change in its secondary structural content from 20 °C to 80 °C, appearing to never appreciably unfold (Figure 6B). After melting, we again collected CD spectra at 80 °C. The CD spectrum of DL116 was largely unchanged, while the DL121 spectrum indicated a loss of secondary structure (Figure 6A). These CD experiments suggest that DL116 is more stable than DL121, but possibly less structurally ordered. Typically, enhanced stability would correspond with enhanced mutational robustness and evolvability. However, DL116 is less mutationally robust and less evolvable in terms of allostery. This implies that the specific mechanism of stabilization in DL116 does not contribute to evolvability in the same way as observed in other proteins (Bloom et al. 2005; Bloom et al. 2006).

**Figure 6:**
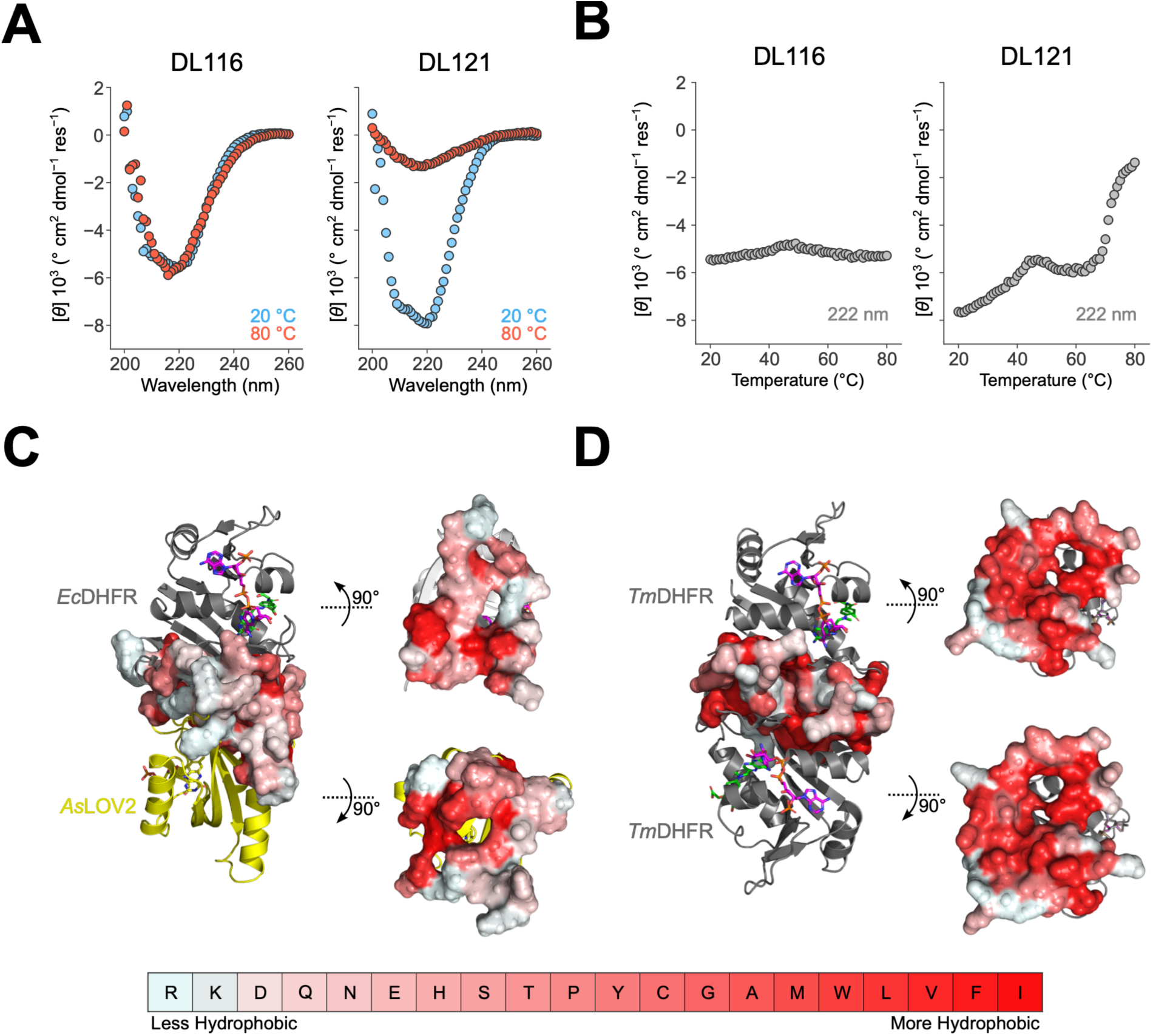
DL116 is a thermostable enzyme. **(A)** Far-UV CD spectra of DL116 and DL121 at 20 °C (blue) and 80 °C (red), following a temperature melt. **(B)** CD temperature melts of DL116 and DL121 measured at 222 nm to monitor changes in alpha helical content. **(C)** An AlphaFold2 model of DL116, with DHFR shown as grey cartoon, LOV2 shown as yellow cartoon, and NADPH, folate, and FMN shown as pink, green, and yellow sticks, respectively. The DHFR residues within 5 Å of LOV2 are shown as a surface colored by the Eisenberg hydrophobicity scale (Eisenberg et al. 1984). Right: isolated DHFR and LOV2 domains are rotated to display interdomain hydrophobicity. **(D)** A crystal structure of the *T. maritima* DHFR dimer (PDB: 1D1G). NADPH and folate are shown as pink and green sticks, respectively. The residues of one DHFR subunit within 5 Å of the other are shown as a surface colored by the Eisenberg hydrophobicity scale (Eisenberg et al. 1984). Right: isolated DHFR subunits are rotated to display interdomain hydrophobicity.

To our knowledge, this is the first example of an engineered domain insertion that results in a hyperstable chimeric protein. To make sense of how insertion of LOV2 at site 115/116 of DHFR increases stability, we compared the AlphaFold model of DL116 to the crystal structure of DHFR from *Thermotoga maritima*, a thermophilic bacterium that grows optimally at 80 °C (Loveridge et al. 2009). *Tm*DHFR (PDB: 1D1G) forms a dimer with an interface that is very similar to the DHFR/LOV2 interface of DL116 (Figure 6C-D). This suggests that the way in which LOV2 docks onto DHFR may be a contributing factor to the enhanced stability of DL116. The interdomain interface of DL116 has a buried surface area of 1,889.89 Å^2^, while the *Tm*DHFR dimer interface is 67% larger with a buried surface area of 3,136.92 Å^2^. Both interfaces also contain buried hydrophobic patches (Figure 6C-D), which may contribute stabilizing interactions between domains. The extensive hydrophobic interface of *Tm*DHFR is known to stabilize the enzyme, at the expense of reduced catalytic efficiency due to constraining the βF-βG loop, whose dynamics are coupled with catalysis (Loveridge et al. 2009). The DHFR/LOV2 interface of DL116 may similarly serve to stabilize the DL116 fusion, while limiting conformational changes that could enable allosteric regulation.

## 3. Discussion

In this study, we asked whether the capacity to evolve allosteric regulation is uniformly distributed across a protein surface or instead depends on local surface context. Using two closely spaced LOV2 insertions in DHFR, we compared the mutational landscapes of allostery when the photoswitch perturbs either a sector-connected insertion site (DL121) or a nearby non-sector-connected insertion site (DL116). Despite similar growth rate phenotypes and enzymatic activity in the dark for the DL121 and DL116 fusions, DL121 exhibited an initial allosteric phenotype and contained multiple significant single mutations that improved the dynamic range of allosteric regulation. In contrast, DL116 lacked measurable regulation *in vitro* or *in vivo* and lacked significant single-mutational paths to gain allosteric regulation. This suggests that, with respect to allosteric regulation, DL116 may be trapped in an evolutionary valley that requires multiple mutations to surmount (Figure 1B). Together, our results support a model in which the evolutionary accessibility of allosteric regulation is influenced by heterogeneous and highly localized properties of a protein surface.

So what features of the protein surface might explain the differential evolutionary accessibility of allostery for DL121 and DL116? One possible explanation is sector connection. While our prior work has shown that sector-connected surfaces can act as hotspots for the initiation of allosteric regulation, these data further indicate that sector connection impacts the evolutionary accessibility of allosteric phenotypes. The rationale is that the sectors identified by SCA represent cooperative networks within proteins that are essential for the propagation of allosteric signals (Süel et al. 2003; Sadovsky and Yifrach 2007). A perturbation at a sector-connected surface would thus be more evolvable because it would not require preliminary mutations to establish cooperative couplings *de novo*. Interestingly, the weak allosteric signals in the DL116 dataset point to the importance of sector connection. Many of the allostery-introducing mutants with p < 0.05 occurred between the end of the Jα helix of LOV2 and the DHFR sector, reminiscent of a “bridge” connecting the Jα helix to the sector. This pattern indicates that any potential routes to achieving allostery, though possibly few in number, may involve the formation of new connections between the site of conformational change and the sector. Together, these findings support the view that protein sectors may play a role in facilitating the evolution of allosteric regulatory sites.

Another potential explanation for the differences in evolutionary accessibility is stability. Our CD measurements show that the secondary structural content of DL116 is remarkably resistant to temperature, implying that it is a highly thermostable enzyme. This runs counter to the prevailing view that protein evolvability is supported by protein stability (Bloom et al. 2005; Bloom et al. 2006). Typically, stability is thought to promote the evolvability of enzymes by reducing the number of mutations that result in unfolding and loss of function, thereby increasing the number of viable evolutionary paths available to the protein. In the case of DL116 and DL121, we found that the opposite was true; DL116 is more stable than DL121, yet has fewer viable mutations, and fewer mutations that introduce an allosteric phenotype. Our results highlight an important nuance in the relationship between protein stability and evolvability, namely that stability may not benefit the evolvability of phenotypes that rely on dynamical properties, such as allosteric regulation. Indeed, prior work suggests that local structural frustration may be a key feature that enables allosteric signaling (Ferreiro et al. 2011).

Our CD data also raises concerns on the initial structural content of LOV2 when inserted into site 115/116 to produce DL116. The CD spectrum of DL116 has a lower mean residual ellipticity at 222 nm than DL121 (Figure 6A). This may suggest a lower alpha helical content in DL116. While we see that the FMN chromophore of DL116 reacts with C450 in response to light (Figure 1D), the Jα and/or A’α helices of DL116 may be constitutively unfolded. This possibility highlights the importance of taking both host and inserted domain tolerance into account when engineered chimeras through domain insertion.

In the domain insertion engineering field, considerable effort has been devoted to identifying sites that both tolerate domain insertion and can support allosteric regulation (Dagliyan et al. 2016). Recently, a language model called ProDomino was created to predict sites that may tolerate domain insertion and possibly elicit allosteric regulation (Wolf et al. 2025). ProDomino scores the C-D loop of *E. coli* DHFR strikingly high compared to the rest of the sequence (Supplementary Figure S8A-B). Notably, analysis of NMR S^2^ order parameters has indicated that the C-D loop of DHFR is highly dynamic in both the E:folate binary complex and the closed E:folate:NADP^+^ ternary complex, unlike the Met20 loop and βF-βG loop, which both have attenuated dynamics in the closed ternary complex (Osborne et al. 2001; Schnell et al. 2004). Previously, some sites in the C-D loop were found to be fairly tolerant to insertion of LOV2 (Reynolds et al. 2011), and the highest-scoring ProDomino sites indeed identify insertion sites that preserve DHFR function. However, quantitative ProDomino prediction of LOV2 insertion tolerance in DHFR was modest, with the model capturing 17-19% of the variance in growth rate data (Supplementary Figure S8C-D). High ProDomino scores were not statistically associated with allosteric effects in DHFR-LOV2 chimeras, explaining only 0.6% of the variance (Supplementary Figure S8E). Because allosteric hotspots in DHFR were previously associated with long-range correlations in sequence information (Reynolds et al. 2011), we investigated whether ProDomino captures similar long-range information in its scoring. We generated insertion tolerance profiles for every DHFR point mutant (Supplementary Figure S9A) and examined the predicted scores at positions 115, 116, 120, and 121 across all mutant backgrounds. Mutations that improved insertion site score were concentrated near the queried positions (Supplementary Figure S9B-E), indicating that local sequence context strongly impacts ProDomino positional scoring. Successful prediction of the evolvability of allosteric regulation may require information describing long-range interactions within proteins.

Together, our data suggest that insertion tolerance alone may be insufficient to predict the evolvability of allosteric regulation. Some sites that have similar domain insertion tolerance, such as the sites used in DL116 and DL121, may differ substantially in their ability to access optimized allostery via single mutational paths. These results are consistent with the idea that a protein’s intrinsic cooperative architecture influences not only where allostery can be introduced in a protein, but also the ease with which allostery can subsequently evolve.

## 4. Methods

### 4.1 Cloning of DL116 into pET24a

The DL116 coding sequence was cloned into a pET24a expression vector with a C-terminal TEV-cleavable hexahistidine tag using restriction-free cloning (rf-cloning.org) (van den Ent and Löwe 2006). A primary PCR contained 10 uL of 5X Q5 buffer, 1 uL of 10 mM dNTPs, 2.5 uL of 10 uM forward and reverse primers, 0.5 uL of Q5 polymerase, and 31.5 uL of nuclease-free water. The reactions were then heated to 98 °C for 30 s, and then cycled through 98 °C for 8 s, 55 °C for 20 s, and 72 °C for 30 s, repeating 35 times with a final heating at 72 °C for 5 min in a Veriti 96-well thermocycler (Applied Biosystems). The primary PCR was run on a 1% agarose gel at 120V for 30 min. The major band containing the synthesized “megaprimer” was excised and purified using a NucleoSpin Gel & PCR Clean-up kit (Macherey-Nagel, ref#740609.50). A secondary PCR contained 4 uL of 5X Q5 buffer, 1 uL of 10 mM dNTPs, 226.8 ng of megaprimer, 73.9 ng of pET24a-DL121, 0.5 uL of Q5 polymerase, and nuclease free water, with a final reaction volume of 20 uL. The secondary PCR was heated to 98 °C for 30 s, and then cycled through 98 °C for 8 s, and then 72 °C for 121 s, repeating 15 times with a final heating at 72 °C for 5 min. Template plasmid was then degraded by adding 2 uL of DpnI to the secondary PCR, and incubating at 37 °C for 2 h. The reaction was then transformed into chemically competent XL1-Blue *E. coli* genotype: recA1 endA1 gyrA96 thi-1 hsdR17 supE44 relA1 lac [F’ proAB lacIqZDM15 Tn10(Tetr)]) (Agilent Technologies).

### 4.2 Purification of DL116 and DL121 proteins

BL21(DE3) *E. coli* (genotype: fhuA2 [lon] ompT gal (l DE3) [dcm] DhsdS. l DE3 = l sBamHIo DEcoRI-B int::(lacI::PlacUV5::T7 gene1) i21 Dnin5) (New England Biolabs) was transformed with a pET24a vector containing either DL116 or DL121. A scraping of colonies was used to inoculate 20 mL of LB, which was incubated overnight at 37 °C with shaking. The following morning, 2 mL of overnight culture was added to 1 L of TB (48 g/L tryptone, 96 g/L yeast extract, 1.6% v/v glycerol, 72 mM K_2_HPO_4_, 17 mM KH_2_PO_4_, 50 μg/mL thymidine, 10 mg/L riboflavin, and 35 μg/mL kanamycin) in a 2,800-mL fernbach flask. The cultures were incubated at 37 °C with shaking at 220 rpm. When the cultures reached an OD_600_ of 0.8, they were placed on ice. DHFR-LOV2 expression was induced by adding 500 uL of 1M IPTG. The induced cultures were incubated overnight at 18 °C. The induced cultures were centrifuged at 5,000 rcf for 20 min at 4 °C, after which the supernatant was discarded and pellets were frozen with liquid nitrogen and stored at −80 °C. Pellets were thawed at 4 °C and resuspended in Nickel Binding Buffer (50 mM Tris, pH 7, 500 mM NaCl, and 10 mM imidazole) supplemented with 100 uM phenylmethylsulfonyl fluoride, 1 μg/mL pepstatin, 10 μg/mL leupeptin, 5 μg/mL DNase, and 100 μg/mL lysozyme, at 5 mL/g of pellet. Cells were then lysed by sonication. The lysate was then fractionated by centrifugation at 40,000 rcf for 30 min at 4 °C. The soluble fraction was collected and incubated with equilibrated Ni-NTA resin (Qiagen, cat#4561) (100 μL slurry/g of cell pellet) for 1 h at 4 °C with continual stirring. The resin was poured into a column and washed twice with 2.5 mL of wash buffer (1M NaCl, 20 mM imidazole, 50 mM tris, pH 7). The his-tagged DHFR-LOV2 protein was then eluted from the column with 3-10 mL of elution buffer (1M NaCl, 400 mM imidazole, 100 mM tris, pH 7) at 4 °C. To cleave the his tag, eluate was mixed with 400-600 uL of purified TEV protease (2.33 mg/mL) and dialyzed against 4 L of TEV cutting buffer (0.5 mM EDTA, 1 mM DTT, 50 mM Tris, pH 7) in 10K MWCO SnakeSkin Dialysis Tubing (Thermo Scientific, ref# 88243) at 4 °C overnight. The protein was again dialyzed against 4 L of TEV binding buffer (1M NaCl, 5% v/v glycerol, 50 mM tris, pH 7) at 4 °C for 4 h. The protein was then diluted to 10 mL with TEV binding buffer. Equilibrated Ni-NTA resin (100 μL slurry/g of cell pellet) was then added to the protein. The mixture was transferred to a column and agitated by pipetting up and down occasionally for 1 hour at 4 °C, after which the flowthrough from the column was collected. The protein was then exchanged into anion exchange buffer A (50 mM tris, pH 7, and 0.0352% v/v β-mercaptoethanol) using a 15-mL 10,000 MWCO centricon. The protein was then concentrated to 1 mL and centrifuged at 22,000 rcf for 10 minutes at 4 °C. The supernatant was then loaded onto a 1-mL anion exchange column (Cytiva, ref#29051325). The protein was then gradient-eluted with anion exchange buffer B (50 mM tris, pH 7, and 0.0352% v/v β-mercaptoethanol, 1M NaCl) using a BioRad NGC Quest FPLC. Fractions containing DHFR-LOV2 were pooled and buffer exchanged into size exclusion chromatography buffer (50 mM tris, pH7, 300 mM NaCl, 1% glycerol) and concentrated to 1 mL using a 4-mL 10,000 MWCO centricon. The protein was then concentrated to 1 mL and centrifuged at 22,000 rcf for 10 minutes at 4 °C. The supernatant was then run on a size exclusion column (HiLoad 16/600 Superdex 75 pg column, GE Life Sciences cat#28989333). Fractions containing DHFR-LOV2 were pooled and concentrated with a 4-mL 10,000 MWCO centricon. The concentrated protein was then frozen with liquid nitrogen and stored at −80 °C.

### 4.3 Steady state Michaelis Menten kinetics of DL116 and DL121

Purified DHFR-LOV2 protein was thawed at 4 °C for 10 minutes and then centrifuged at 22,000 rcf for 10 minutes at 4 °C. The supernatant was then transferred to a new tube. The protein was then diluted to 10 μM in MTEN buffer (50 mM MES, 25 mM tris, pH 7, 25 mM ethanolamine, 100 mM NaCl), supplemented with 5 mM DTT. The concentration of the diluted protein was then quantified five times by absorbance at 280 nm (ε = 44,920 M^−1^ cm^−1^) using a Nanodrop and averaged. The protein was then diluted to 110 nM in MTEN buffer with 5 mM DTT and 100 μM NADPH. Solutions of DHF were prepared in MTEN buffer with 5 mM DTT at 10X concentrations (1 mM, 500 μM, 250 μM, 100 μM, 50 μM, 25 μM, 10 μM, and 5 μM). Aliquots of protein solution and DHF solution were kept on ice before transferring to a 17 °C water bath. DL121 was previously shown to have a maximal allosteric effect at 17 °C (Lee et al. 2008). After 3 minutes in the water bath, 900 uL of protein was mixed with 100 uL of DHF solution. The mixture was then transferred to a quartz cuvette with a 1 cm path length and the absorbance at 340 nm was monitored for 1 min in a Lambda 650 UV/Vis spectrophotometer with a Peltier system set to 17 °C. Lit samples were illuminated by a full spectrum 125 watt 6400K compact fluorescent bulb (Hydrofarm Inc., cat#FLC125D) mounted about 25 cm from the samples during the 3 min incubation in the waterbath, and for 7 s while in a pipette tip, held about 15 cm from the lamp. Dark samples were exposed to the lamp during the 3 min incubation in the water bath, but were kept in opaque tubes. Mixing of dark samples was performed with the lamp turned off in a dark room. Initial velocities, *V_o_*,were calculated using linear regression over the first 15 s of absorbance data. The Michaelis-Menten equation, 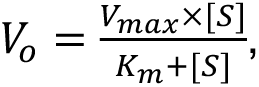 was fit using the SciPy optimize library.

### 4.4 Spectral scan of the LOV2 domain

Purified DHFR-LOV2 protein was thawed at 4 °C for 10 minutes and then centrifuged at 22,000 rcf for 10 minutes at 4 °C. The supernatant was then transferred to a new tube. The protein was diluted to 50 μM in MTEN buffer with 5 mM DTT. The protein was illuminated by a full spectrum 125-watt 6400K compact fluorescent bulb (Hydrofarm Inc., cat#FLC125D) to bring the protein into the lit state. The protein was then loaded into a quartz cuvette with a 1 cm path length. Its absorbance spectrum was then immediately measured from 350 nm to 550 nm using a Lambda 650 UV/Vis spectrophotometer with a Peltier system set to 17 °C. The protein was allowed to relax into the dark state while inside the spectrophotometer for 10 min. The dark state absorbance spectrum was then measured.

### 4.5 Saturation mutagenesis of DL116

The saturation mutagenesis library of the DHFR domain of DL116 was constructed as described in Thompson et al. 2020 (Thompson et al. 2020). Four sublibraries were constructed in which point mutants were contained within one of four regions of the DHFR sequence: positions 1-40 (sublibrary 1, SL1), positions 41-80 (sublibrary 2, SL2), positions 81-115 (sublibrary 3, SL3), and positions 116-159 (sublibrary 4, SL4). Inverse PCR was performed using primers that mutate each sense codon in the DHFR sequence of DL116 to NNS codons, thus producing all possible amino acid substitutions. The NNS mutagenic primers were first phosphorylated using T4 polynucleotide kinase (PNK; NEB, cat#M0201S). Each 20 μL phosphorylation reaction contained 16.5 μL sterile water, 2 μL T4 ligase buffer, 0.5 μL T4 PNK, and 1 μL of 100 μM NNS primer. Reactions were incubated at 37 °C for 1 h, followed by enzyme inactivation at 65 °C for 20 min. PCR amplification was carried out using 2x Q5 Master Mix (NEB, cat#M0492), 10 ng plasmid template, and 500 nM each of forward and reverse primers. Thermocycling conditions consisted of an initial denaturation at 98 °C for 30 s, followed by 22 cycles of 98 °C for 10 s, 55 °C for 30 s, and 72 °C for 2 min, with a final extension at 72 °C for 5 min. To remove methylated template DNA, 1 μL DpnI (NEB, cat#R0176) was added to 25 μL of PCR product and incubated at 37 °C for 4 h. Amplified DNA was purified by gel extraction followed by cleanup with a DNA Clean and Concentrator-5 kit (Zymo Research, cat#D4014).

Purified PCR products underwent a second phosphorylation step containing 100 μL gel-purified DNA, 12 μL 10x T4 ligase buffer, 5 μL T4 PNK, and 5 μL sterile water. Reactions were incubated at 37 °C for 1 h and heated to 90 °C for 30 s. Ligation reactions were assembled using 100 μL of phosphorylated PCR product, 15 μL T4 DNA ligase (NEB, cat#M0202S), 30 μL T4 ligase buffer, and 155 μL sterile water, and were incubated at room temperature for 24 h. Product concentrations were determined by gel densitometry using ImageJ, and individual reactions were pooled in equimolar amounts to generate sublibraries. Four sublibraries were constructed, spanning residues 1–40 (sublibrary 1), 41–80 (sublibrary 2), 81–120 (sublibrary 3), and 121–150 (sublibrary 4). Each sublibrary was transformed into electrocompetent XL1-Blue E. coli using a MicroPulser Electroporator (Bio-Rad) and Gene Pulser cuvettes (Bio-Rad, cat#165-2089). Plasmids were isolated using a GeneJET Plasmid Miniprep Kit (Thermo Scientific, cat#K05053), and library coverage was confirmed by deep sequencing on an Illumina MiSeq platform.

### 4.6 Continuous culture of DL116

A 4-mL culture of *E. coli* ER2566 Δ*folA* Δ*thyA* was inoculated from a frozen glycerol stock into 4 mL of LB media supplemented with 50 μg/mL of thymidine and grown overnight at 37 °C with shaking. The following morning, 200 μL of culture was added to 20 mL of LB media supplemented with 50 μg/mL of thymidine and grown at 37 °C with shaking until the OD_600_ reached 0.4-0.6. The culture was then put on ice and washed three times by centrifuging at 6,500 rcf for 1.5 min at 4 °C, and resuspending in chilled 10% v/v glycerol. One hundred micrograms of each sublibrary was then individually transformed by electroporation using a MicroPulser Electroporator (Bio Rad) and gene pulser cuvettes (Bio Rad, cat#165–2089), and recovered in 1.2 mL of SOB media. Serial dilutions of the electroporated library were plated to check transformation efficiency, and the rest of the outgrowth was spun down at 3,500 rcf for 5 min. The supernatant was discarded and the pellet was resuspended in 20 mL of M9 media supplemented with products of the folate metabolic pathway (93.0 mM sodium (Na^+^), 22.1 mM potassium (K^+^), 18.7 mM ammonium (NH_4_) with 0.4% glucose, 50 μg/mL of thymidine, 22 μg/mL of adenosine, 1 μg/mL of calcium pantothenate, 38 μg/mL of glycine, and 37.25 μg/mL of methionine, and 30 μg/mL of chloramphenicol). The library was then grown overnight in a 125-mL flask at 37 °C with shaking. The culture was then diluted to an OD_600_ of 0.1 in 200 mL of M9 media with 0.4% glucose, 50 μg/mL of thymidine, and 30 μg/mL of chloramphenicol. The cultures were then adapted in 1-L flasks at 30 °C with shaking for 4 h. The adapted cultures were then washed three times by centrifuging at 2,000 rcf for 10 min and resuspending the pellet in M9 media with 0.4% glucose, 1 μg/mL of thymidine, and 30 μg/mL of chloramphenicol. The cultures were then diluted to an OD_600_ of 0.1 in M9 media with 0.4% glucose, 1 μg/mL of thymidine, and 30 μg/mL of chloramphenicol. Sixteen milliliters of diluted culture was then dispensed into glass turbidostat vials. The turbidostat design was based on that described in Toprak et al. 2013, and was outfitted with blue LEDs as in McCormick et al. 2021 (Toprak et al. 2013; McCormick et al. 2021). The OD_600_ was continuously monitored throughout the experiment, and the turbidostat was set to clamp at an OD_600_ of 0.15. The turbidostat was kept at 30 °C, and each vial was continuously stirred with a stir bar. Vials designated as ‘lit’ were each illuminated with a single 5V blue LED throughout the experiment. There were three replicate ‘lit’ and ‘dark’ vials. One milliliter samples were taken from each vial at the beginning of selection (t = 0 h), and after 4, 8, 12, 16, 20, and 24 h. The samples were centrifuged at 21,130 rcf for 5 min at room temperature and the pellet was stored at −20 °C.

### 4.7 Deep sequencing of DL116 samples from continuous culture

Samples taken from the turbidostat were lysed by resuspending the pellets in 10 μL of sterile water and heating on a 95 °C heat block for 5 min. TruSeq adapters were then added onto the mutated region of each sublibrary by PCR. One microliter of lysate was mixed with 13.25 μL of nuclease-free water, 5 μL of Q5 buffer, 0.5 μL of 10 mM dNTPs, 2.5 μL each of 10 μM TruSeq forward and reverse primer, and 0.25 μL of Q5 polymerase. The reactions were held at 98 °C for 90 s, then cycled 20 times through 98 °C for 10 s, 63 °C for SL1 and SL2 or 64 °C for SL3 and SL4 for 15 s, and 72 °C for 10 s, with a final extension at 72 °C for 120 s in a Veriti 96-well thermocycler (Applied Biosystems). Unique i5/i7 indexing primers were then used for each sample in a secondary PCR: 1 μL of each PCR product was combined with 13.25 μL of nuclease-free water, 5 μL of Q5 buffer, 0.5 μL of 10 mM dNTPs, 2.5 μL each of 10 μM forward and reverse primer, and 0.25 μM Q5 polymerase. The secondary PCRs were held at 98 °C for 90 s, then cycled 20 times through 98 °C for 10 s, 64 °C for 15 s, and 72 °C for 10 s, with a final extension at 72 °C for 120 s. The concentration of double-stranded DNA in each reaction was quantified using a PicoGreen Assay (Thermo Scientific, cat#P7589) on a Victor X3 multimode plate reader (Perkin Elmer). One microgram of dsDNA from each reaction was then mixed and purified by gel extraction using a NucleoSpin Gel & PCR Clean-up kit (Macherey-Nagel, ref#740609.50). DNA purity was determined by measuring the 260 nm/230 nm and 260 nm/280 nm absorbance ratios using a DS-11 spectrophotometer (DeNovix). DNA concentration was determined using the Qubit 3 (Thermo Scientific). The pooled amplicon library was then sent to Azenta Life Sciences where it was analyzed by TapeStation (Agilent Technologies) and sequenced with Illumina sequencing (Genewiz) with a 2 x 150 bp dual index run with 20% PhiX spike-in yielding 725,852,554 reads. We randomly downsampled DL116 reads and DL121 reads from McCormick et al. 2021 to achieve a contribution of 637,446,766 reads from each dataset that were used in our subsequent analyses.

### 4.8 Read joining, downsampling, filtering, and counting

The forward and reverse reads provided by Azenta were joined using usearch v11.0.667 using the i86linux32 package. The T0 reads for DL116 were randomly downsampled to be equal to the T0 reads from DL121. The mean DL116 reads across subsequent timepoint samples was then calculated. DL121 reads were randomly downsampled to the mean of DL116. The sublibrary identity of each read was identified using substring searches. The reads were then trimmed to the coding region of each sublibrary and filtered such that a read was discarded if any nucleotide in its coding region had a Q score below 30. Mutations were then identified and counted in each of the samples. Illumina sequencing error rates were estimated using the “Hamming Correction” method described in McCormick et al. 2021 (McCormick et al. 2021). The estimated number of errant reads (attributed to sequencing noise around the unmutated construct) was subtracted from the mutant and wild-type codon-level counts.

### 4.9 Computing relative growth rate and allosteric effect size

The Hamming-corrected counts were then used to compute relative mutant frequency as a function of time according to the following equation:

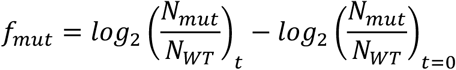

The growth rates of each mutant in the light and dark were calculated using weighted least squares linear regression of *f_mut_* over time, weighted by the number of hamming-corrected counts for that mutant in the sample. The growth rate (g) was then computed with weighted least squares linear regression of log_2_(g) vs. time. The linear regression was weighted with sequencing counts to mitigate the influence of counting noise from data points with few counts (Rubin et al. 2017). The “allostery tuning score” (ΔA) of each mutant was then calculated by subtracting the mean dark growth rate from the mean lit growth rate. Since the allostery tuning scores are relative, the allostery-tuning scores of unmutated DL116 and DL121 are both equal to zero. However, the allosteric effect size of unmutated DL116 and DL121 differ. To improve interpretability, we added the baseline allosteric effect (A_0_) to each of the allostery tuning scores to calculate the allosteric dynamic range (ADR) of each mutant (see Supplementary Figure S7).

### 4.10 Statistical analyses

Prior to assessing significant allosteric mutations, we performed data quality filtering. The average growth rate of stop codons in the first two sublibraries (positions 1-80 of DHFR) was used as a viability cutoff. Mutants with a computed growth rate below this floor were not considered in the analysis. Additionally, we only considered mutants with at least 50 counts at timepoint zero and mutants that were present in at least five of the seven timepoints. P-values for the allosteric effect sizes were calculated using a Welch’s t-test under the null hypothesis that the lit and dark growth rates have equal means. For each chimera, two significance cutoffs were used to determine significant allostery-tuning mutants. The standard cutoff of p=0.05 and cutoffs determined using Sequential Goodness of Fit (SGoF) of p = 0.0089 for DL116 and p = 0.0083 for DL121 (Carvajal-Rodriguez and Uña-Alvarez 2011). To compare the overall enrichment of significant allosteric mutants in DL116 and DL121, we performed a label-permutation enrichment test. After filtering mutants for viability and measurement quality, the chimera identity labels for each mutant were removed and their p-values were pooled. P-values were then randomly re-allocated to either DL116 or DL121 equal to the number of viable mutants for each. The number of p-values below 0.05 were counted for each random re-allocation to make distributions of the number of significant mutants expected by random chance. These null distributions were then compared to the observed number of mutants with p < 0.05 for each chimera.

### 4.11 Circular dichroism spectroscopy

Purified DHFR-LOV2 protein was thawed at 4 °C for 10 min and exchanged into CD buffer (50 mM sodium phosphate, pH 6.5, 100 mM sodium fluoride) using a Pierce Protein Concentrator PES 10K MWCO (Thermo ref#88517). The samples were then centrifuged at 22,000 rcf at 4 °C for 5 min. No pellet was observed. The sample was transferred to a new tube and the protein concentration was measured five times by absorbance at 280 nm (ε = 44,920 M^−1^ cm^−1^) using a Nanodrop and averaged. The protein was then diluted to 0.75 mg/mL in CD buffer. The sample was then loaded into a cuvette and placed in an AVIV Model 420 Circular Dichroism Spectrometer, where it was allowed to relax into the dark state at 20.00 °C. A spectral scan was then collected at 20.00 °C from 260.00 to 195.00 nm with a bandwidth of 1.00 nm, an averaging time of 3.000 s, and a settling time of 0.330 s. The sample was then heated from 20.00 °C to 80.00 °C with a 1.00 °C/min ramp rate and a 1.00 min temperature equilibration time. The CD signal at 220.00 nm was monitored over the course of the melt with a bandwidth of 1.00 nm and an averaging time of 3.00 s. Following the melt, another spectral scan was collected at 80.00 °C. The sample was visually inspected after data acquisition, at which point it was observed that DL121 formed aggregates, whereas DL116 remained in solution.

### 4.12 Prediction of Domain Insertion Tolerance with ProDomino

ProDomino code was cloned from https://github.com/Niopek-Lab/ProDomino and run on a Google Colab Pro server (T4 GPU and high RAM) with one minor change: a function to convert 3-letter amino acid code to 1-letter amino acid code was imported from a newer Biopython module under a wrapper. The ProDomino embedder and model were used to generate insertion site prediction scores for every residue in the wild-type *E. coli* DHFR sequence. These scores were then compared to growth rate data for DHFR-LOV2 insertions from Reynolds et al. 2011 (Reynolds et al. 2011). The ProDomino embedder and model were then used to generate insertion prediction scores for all sequences corresponding to site-saturation mutagenesis of the *E. coli* DHFR sequence, and predicted scores were compared to experimental data.

## Supporting information

Supplementary Table 2 - growth rate data

## Acknowledgements

We would like to thank members of the Reynolds lab and Scott Saunders’s lab at UT Southwestern Medical Center (UTSW) who provided feedback on this work. We would also like to thank Chad Brautigam and Scott Tso from the Macromolecular Biophysics Resource at UTSW for help with early CD experiments on DL116. We would like to thank Katie Tripp at the Center for Molecular Biophysics at Johns Hopkins University for help with CD spectroscopy. Analysis of NGS data was performed on the UT Southwestern BioHPC. Research was supported by grants T32GM131963 to JD, T32GM141804 to RS, and NSF Grant #1942354 to KR.

## Author Contributions

Conceptualization: KR. Investigation: JD, JM, RS, ST. Methodology: JM. Formal analysis: JD. Writing — Original Draft: JD, KR. Writing — Review & Editing: JD, JM, RS, ST, KR. Supervision: KR. Funding Acquisition: KR.

## Code and Data Availability

Code to replicate all analyses can be found on GitHub: https://github.com/reynoldsk/evolvability_DLfusions. Illumina sequencing data for DL116 has been deposited in the NCBI SRA under BioProject: PRJNA1514100. The DL121 Illumina sequencing data was previously deposited under BioProject: PRJNA706683.

