## Supplementary Materials for "Protein Surface Site Determines the Evolutionary Accessibility of Allosteric Regulation"

**Contents:**


**I. Supplementary Figures**

**Figure S1:** FASTQ Downsampling.

**Figure S2:** Library compositions at experiment start.

**Figure S3:** Heatmaps of growth rates.

**Figure S4:** Correlations of DL121 mutant growth rates and allostery with values from McCormick et al. 2021.

**Figure S5:** Inter-replicate growth rate reproducibility.

**Figure S6:** Enrichment of deleterious mutants in the DHFR sector.

**Figure S7:** Computing allosteric effects in DL116 and DL121.

**Figure S8:** Comparison of DHFR-LOV2 insertion scan from Reynolds et al. 2011 to ProDomino predictions of domain insertion tolerance.

**Figure S9:** ProDomino predictions for DHFR single mutants.

**II. Supplementary Tables**

**Table S1:** Kinetic Parameters and growth rates of unmutated DL116 and DL121

**Table S2:** Relative growth rates of DL116 and DL121 chimeras **(provided as a .csv file)**

**A**

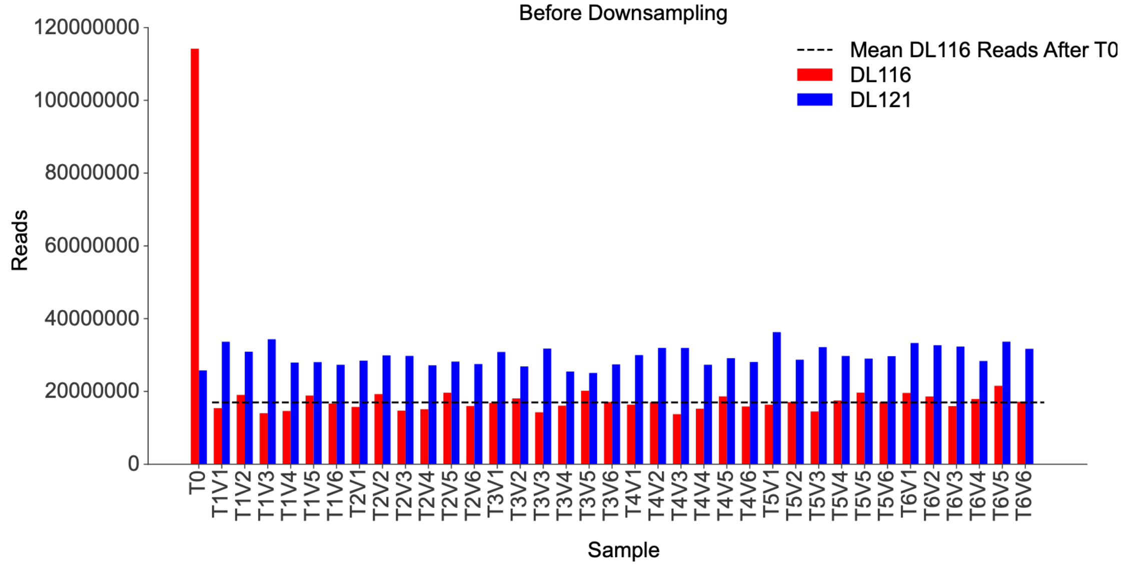


**B**

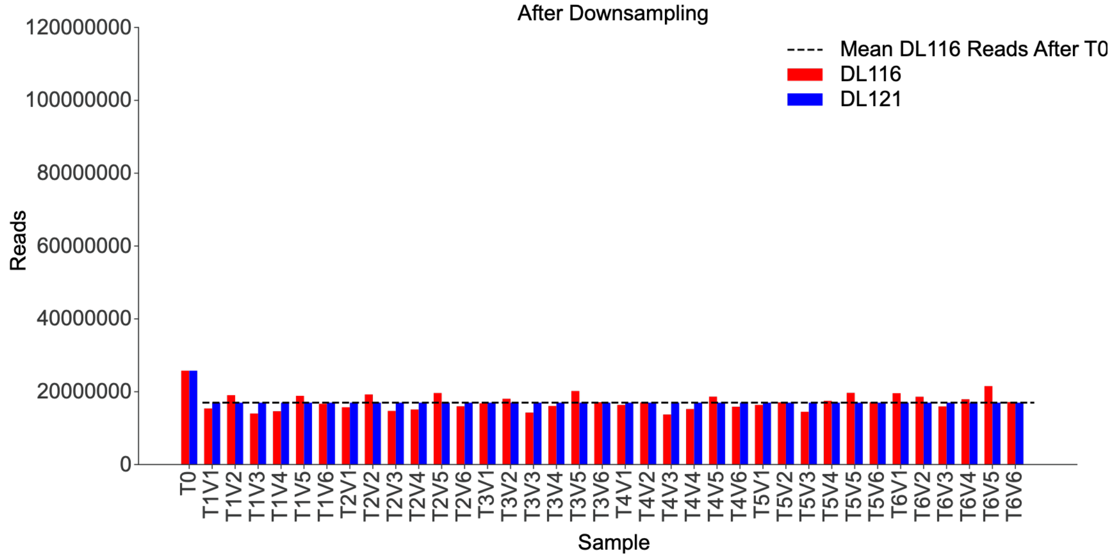


**Supplementary Figure S1: FASTQ Downsampling. (A)** Barplot of the number of reads assigned to each turbidostat sample before downsampling. **(B)** Barplot of the number of reads assigned to each turbidostat sample after downsampling.

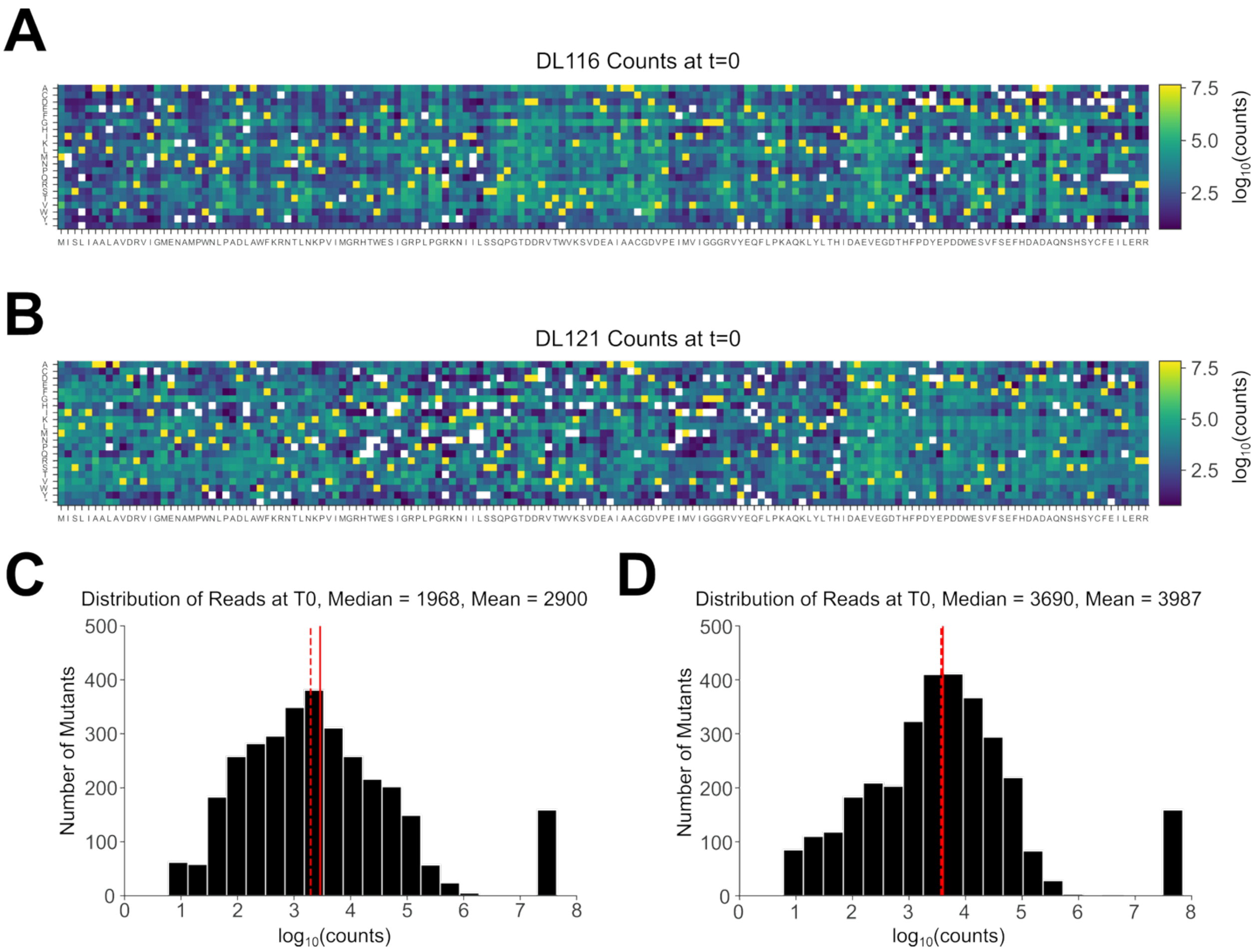


**Supplementary Figure S2: Library compositions at experiment start. (A-B)** Heatmaps depicting the $\log_{10}$(counts) at the start of the experiment for DL116 (A) and DL121 (B). The horizontal axes are the wild-type sequence of DHFR. The vertical axes are the possible amino acid substitutions. **(C-D)** Histograms of $\log_{10}$(counts) at the start of the experiment for DL116 (C) and DL121 (D). The solid and dashed red lines are the mean and median, respectively.

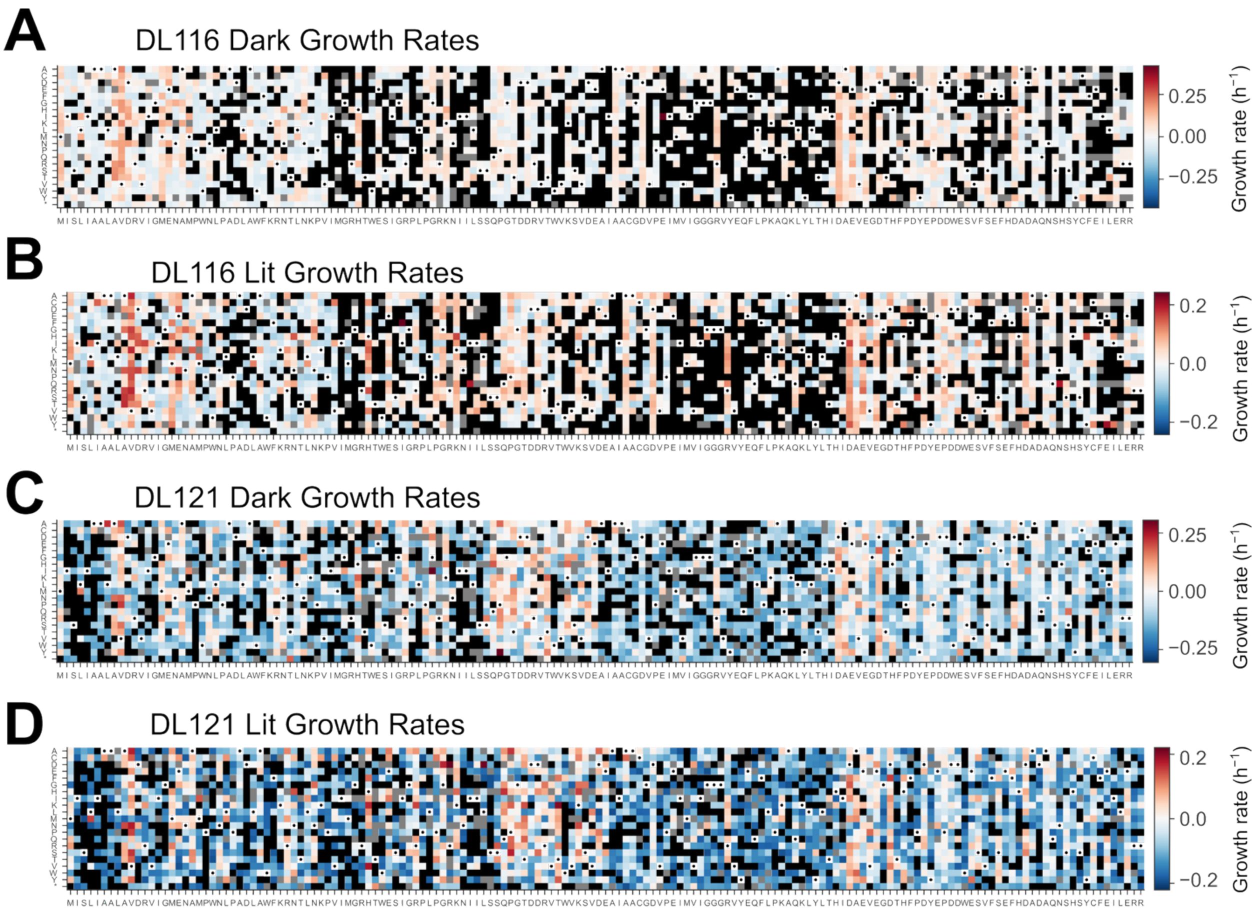


**Supplementary Figure S3: Heatmaps of growth rates. (A-D)** Heatmaps of mutant growth rates relative to the unmutated chimeras for (A) DL116 in the dark, (B) DL116 in the light, (C) DL121 in the dark, (D) DL121 in the light. Mutants that increased relative growth rate are shown in red, whereas mutants that decreased relative growth rate are shown in blue. Mutants that did not affect growth rate are shown in white. Pixels corresponding to wild-type identity contain a black circle. Mutants with growth rates lower than the mean of stop codons in the first 80 positions of DHFR are nonviable and not reliably measured and shown as black pixels. Mutants that had insufficient data to fit a growth rate are shown as grey pixels.

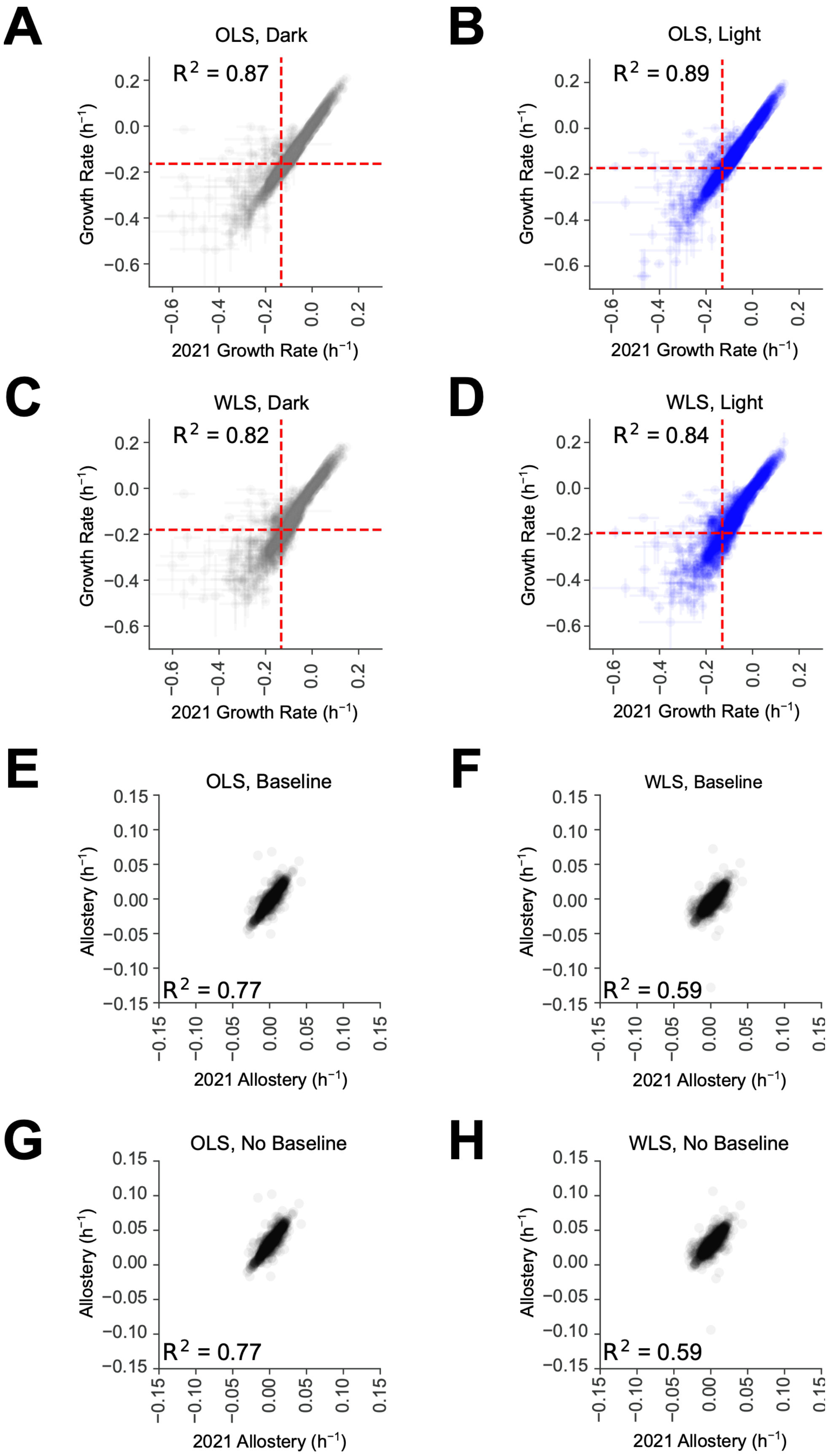
A
OLS, Dark
$R^2 = 0.87$
Growth Rate ($h^{-1}$)
2021 Growth Rate ($h^{-1}$)
B
OLS, Light
$R^2 = 0.89$
Growth Rate ($h^{-1}$)
2021 Growth Rate ($h^{-1}$)
C
WLS, Dark
$R^2 = 0.82$
Growth Rate ($h^{-1}$)
2021 Growth Rate ($h^{-1}$)
D
WLS, Light
$R^2 = 0.84$
Growth Rate ($h^{-1}$)
2021 Growth Rate ($h^{-1}$)
E
OLS, Baseline
$R^2 = 0.77$
Allostery ($h^{-1}$)
2021 Allostery ($h^{-1}$)
F
WLS, Baseline
$R^2 = 0.59$
Allostery ($h^{-1}$)
2021 Allostery ($h^{-1}$)
G
OLS, No Baseline
$R^2 = 0.77$
Allostery ($h^{-1}$)
2021 Allostery ($h^{-1}$)
H
WLS, No Baseline
$R^2 = 0.59$
Allostery ($h^{-1}$)
2021 Allostery ($h^{-1}$)

**Supplementary Figure S4: Correlations of DL121 mutant growth rates and allostery with values from McCormick et al. 2021. (A-D)** Correlations of DL121 mutant growth rates determined using ordinary least squares (OLS) (A-B) or weighted least squares (WLS) (C-D) linear regression and DL121 mutant growth rates from McCormick et al. 2021. Circles represent the average of three replicate measurements and error bars represent the standard deviation. The horizontal red dashed lines depict the average growth rate of nonsense mutations in positions 1-80 of DHFR. The vertical red dashed lines depict the growth rate of DL121-D27N, which was used as a growth rate cutoff in McCormick et al. 2021. **(E-F)** Correlations of allosteric dynamic range (ADR), calculated as the allostery-tuning effect (ΔA) plus the initial, or baseline, allostery ($A_0$), derived from growth rates determined with either OLS (E) or WLS (F) linear regression. **(G-H)** Correlations of allostery-tuning effects (ΔA) to those from McCormick et al. 2021, calculated as the difference in relative growth rate in the light and dark ($g_{lit} - g_{dark}$) determined from with either OLS (G) or WLS (H) linear regression.

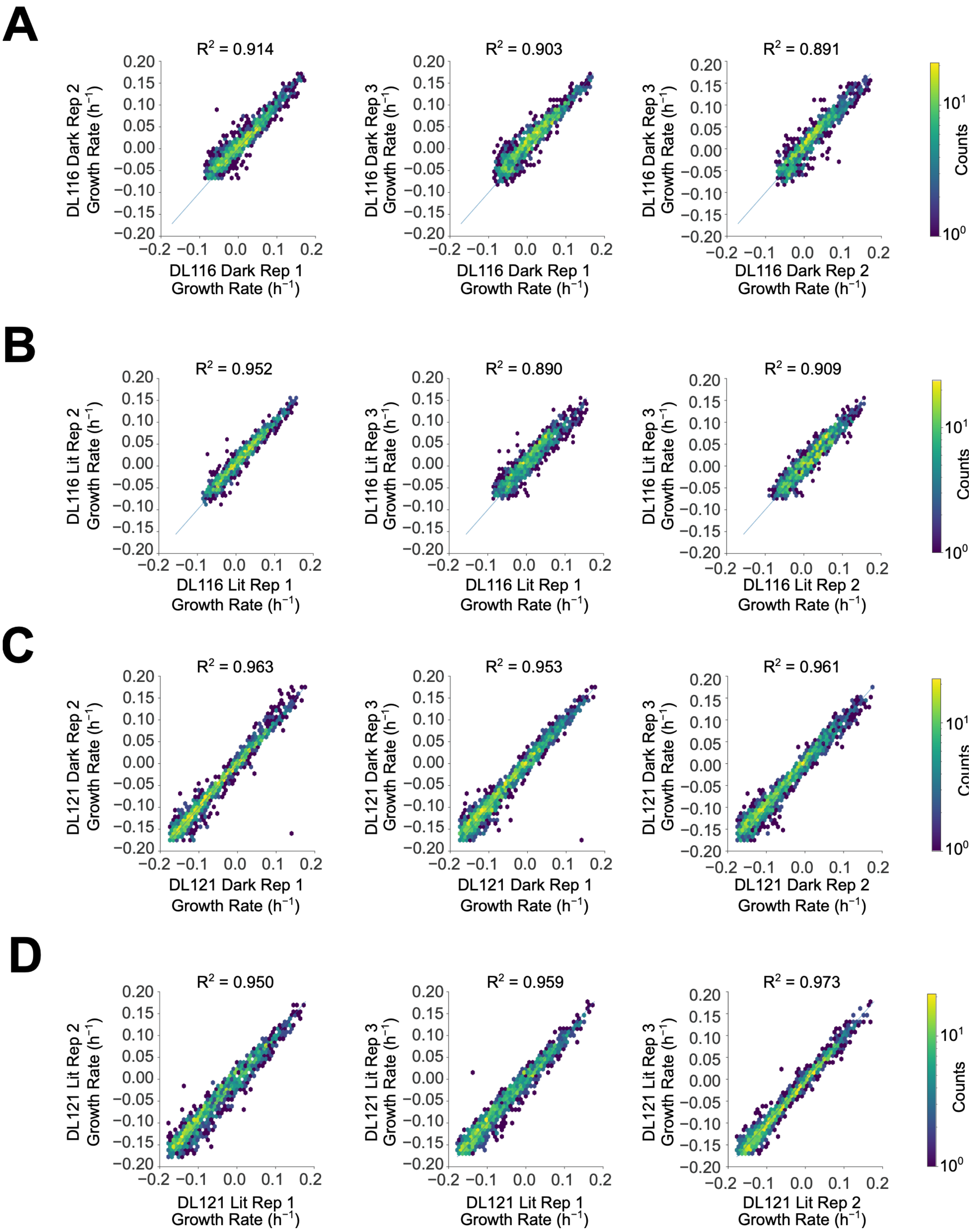

A
R2 = 0.914
R2 = 0.903
R2 = 0.891
DL116 Dark Rep 2 Growth Rate (h−1)
DL116 Dark Rep 3 Growth Rate (h−1)
DL116 Dark Rep 1 Growth Rate (h−1)
DL116 Dark Rep 2 Growth Rate (h−1)
Counts
B
R2 = 0.952
R2 = 0.890
R2 = 0.909
DL116 Lit Rep 2 Growth Rate (h−1)
DL116 Lit Rep 3 Growth Rate (h−1)
DL116 Lit Rep 1 Growth Rate (h−1)
DL116 Lit Rep 2 Growth Rate (h−1)
Counts
C
R2 = 0.963
R2 = 0.953
R2 = 0.961
DL121 Dark Rep 2 Growth Rate (h−1)
DL121 Dark Rep 3 Growth Rate (h−1)
DL121 Dark Rep 1 Growth Rate (h−1)
DL121 Dark Rep 2 Growth Rate (h−1)
Counts
D
R2 = 0.950
R2 = 0.959
R2 = 0.973
DL121 Lit Rep 2 Growth Rate (h−1)
DL121 Lit Rep 3 Growth Rate (h−1)
DL121 Lit Rep 1 Growth Rate (h−1)
DL121 Lit Rep 2 Growth Rate (h−1)
Counts

**Supplementary Figure S5: Inter-replicate growth rate reproducibility. (A-D)** Two-dimensional hex bin histograms correlating growth rates computed for different replicates of (A) DL116 in the dark, (B) DL116 in the light, (C) DL121 in the dark, (D) DL121 in the light. Data shown are those with growth rates above the mean growth rate for nonsense mutations at positions 1-80 of DHFR in matched conditions ("viable mutants"). Coefficients of determination ($R^2$) were computed for viable mutants.

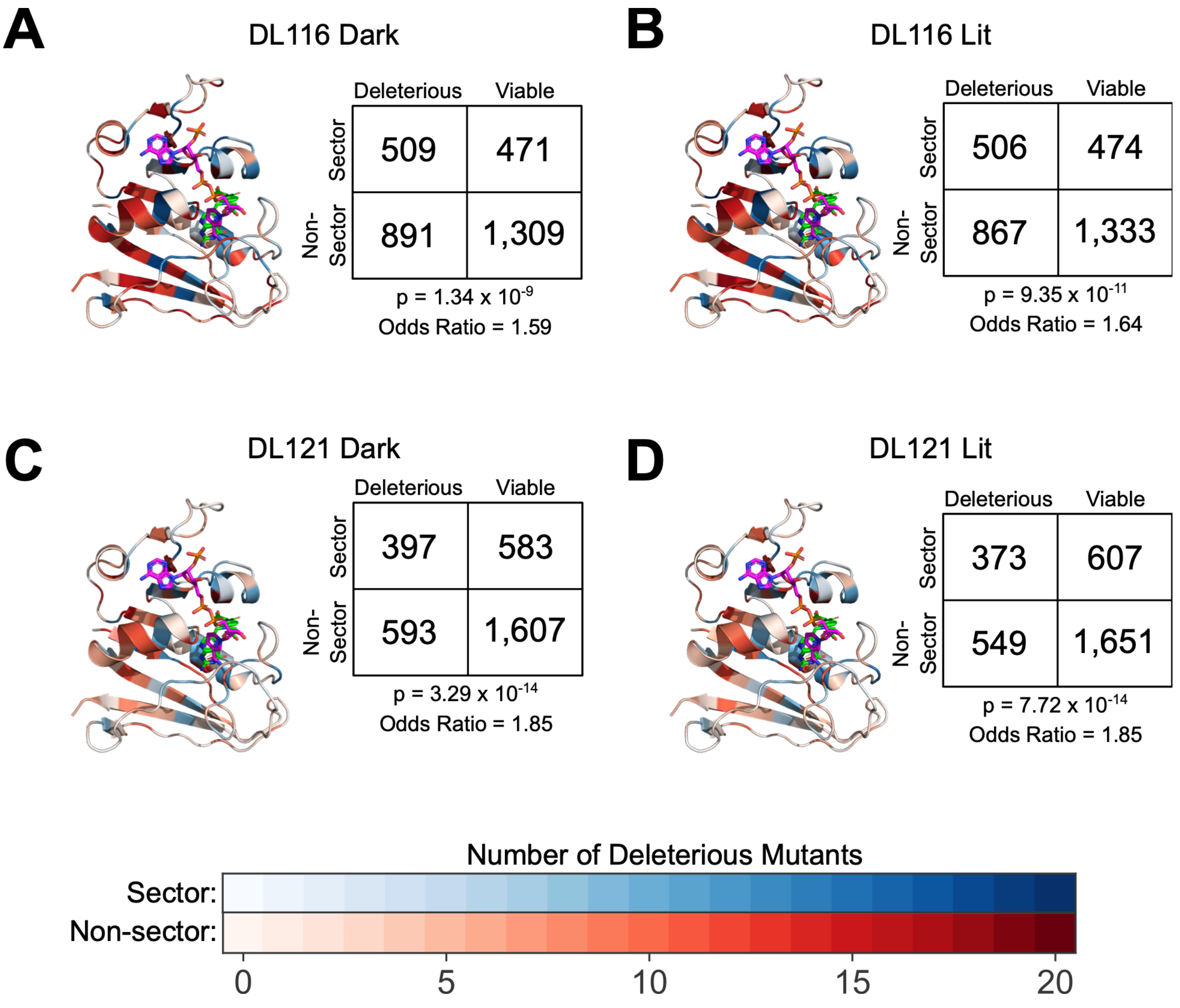


**Supplementary Figure S6: Enrichment of deleterious mutants in the DHFR sector. (A)** DL116 in the dark, **(B)** DL116 in the light, **(C)** DL121 in the dark, **(D)** DL121 in the light. In each panel, the structure of DHFR (PDB: 1RX2) is colored by sector identity and the number of deleterious mutants across positions. Deleterious is defined as having a growth rate less than or equal to the mean growth rate of nonsense mutations in the first 80 positions of DHFR, or having insufficient data to fit growth rates (these mutants are assumed to be deleterious). Contingency tables are shown, along with the results of Fisher's exact tests. Deleterious mutants are significantly enriched in the sector in both DL116 and DL121, in both the light and dark. NADPH and folate are shown as pink and green sticks, respectively.

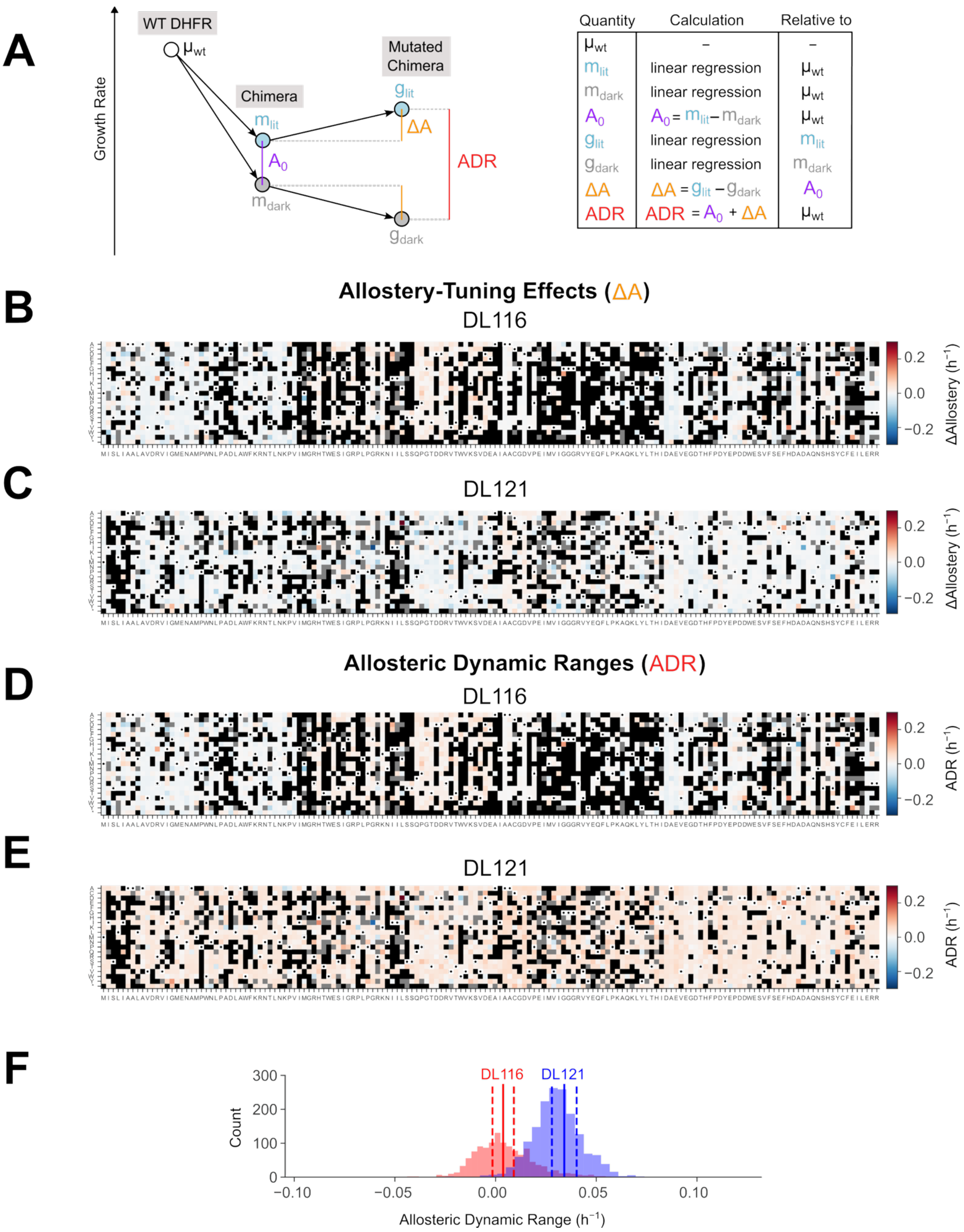
A
WT DHFR
μwt
Chimera
Mutated Chimera
Growth Rate
mlit
mdark
glit
gdark
A0
ΔA
ADR
Quantity
Calculation
Relative to
μwt – –
mlit linear regression μwt
mdark linear regression μwt
A0 A0 = mlit – mdark μwt
glit linear regression mlit
gdark linear regression mdark
ΔA ΔA = glit – gdark A0
ADR ADR = A0 + ΔA μwt
B
Allostery-Tuning Effects (ΔA)
DL116
ΔAllostery (h⁻¹)
0.2
0.0
−0.2
C
DL121
ΔAllostery (h⁻¹)
0.2
0.0
−0.2
D
Allosteric Dynamic Ranges (ADR)
DL116
ADR (h⁻¹)
0.2
0.0
−0.2
E
DL121
ADR (h⁻¹)
0.2
0.0
−0.2
F
DL116
DL121
Count
300
200
100
0
−0.10
−0.05
0.00
0.05
0.10
Allosteric Dynamic Range (h⁻¹)

**Supplementary Figure S7: Computing allosteric effects in DL116 and DL121. (A)** Explanation of growth rate and allostery calculations. Left: The growth rate corresponding to wild-type DHFR ($\mu_{wt}$) is represented along a vertical growth rate axis. LOV2 insertion reduces the growth rate and produces two new relative growth rates ($m_{lit}$ and $m_{dark}$), which differ by $A_0$, the initial baseline allosteric effect of that DHFR-LOV2 chimera. The growth rates of DHFR-LOV2 mutants ($g_{lit}$ and $g_{dark}$) are calculated relative to $m_{lit}$ and $m_{dark}$. The difference between $g_{lit}$ and $g_{dark}$, $\Delta A$, describes the change in allosteric effect size from the unmutated chimera. The absolute allosteric dynamic range (ADR) of each mutant is calculated by summing the initial allostery $A_0$ and the allostery tuning effect $\Delta A$, in a process we refer to as "baseline correction." Right: A table summarizing how each value is calculated, and which value it is relative to. **(B-C)** Heatmaps of allostery-tuning scores ($\Delta A$) relative to unmutated (B) DL116 or (C) DL121. These scores are computed as the difference in relative growth rate in the light and dark ($m_{lit}$ - $m_{dark}$) and reflect how much the allosteric effect size has changed as a result of mutation. In DL116 (B), red pixels are mutants that introduce positive allostery, blue pixels introduce negative allostery, and white pixels do not introduce allostery. In DL121 (C), red pixels are mutants that enhance existing allostery, blue pixels are mutants that disrupt allostery, and white pixels do not affect allostery. **(D-E)** Heatmaps of allosteric dynamic range (ADR) for mutants of (D) DL116 or (E) DL121. These scores are equal to the allostery tuning scores from panels B and C, plus the initial dynamic range in growth rate of the unmutated chimera relative to WT DHFR. These scores reflect the overall dynamic range of allostery observed in the resulting mutants. Red pixels are mutants that have positive allostery, while blue pixels are mutants with negative allostery. White pixels are mutants with no allostery. Pixels corresponding to wild-type identity are white and contain a black circle. Mutants with growth rates lower than the mean of stop codons in the first 80 positions of DHFR are deemed nonviable and be unreliably measured and are shown as black pixels. Mutants that had insufficient data to fit a growth rate are shown as grey pixels. **(F)** Distributions of the allosteric dynamic range (ADR) of DL116 (red) and DL121 (blue) mutants. The solid vertical lines indicate the mean, and the dashed lines indicate the mean ± standard error of the mean.

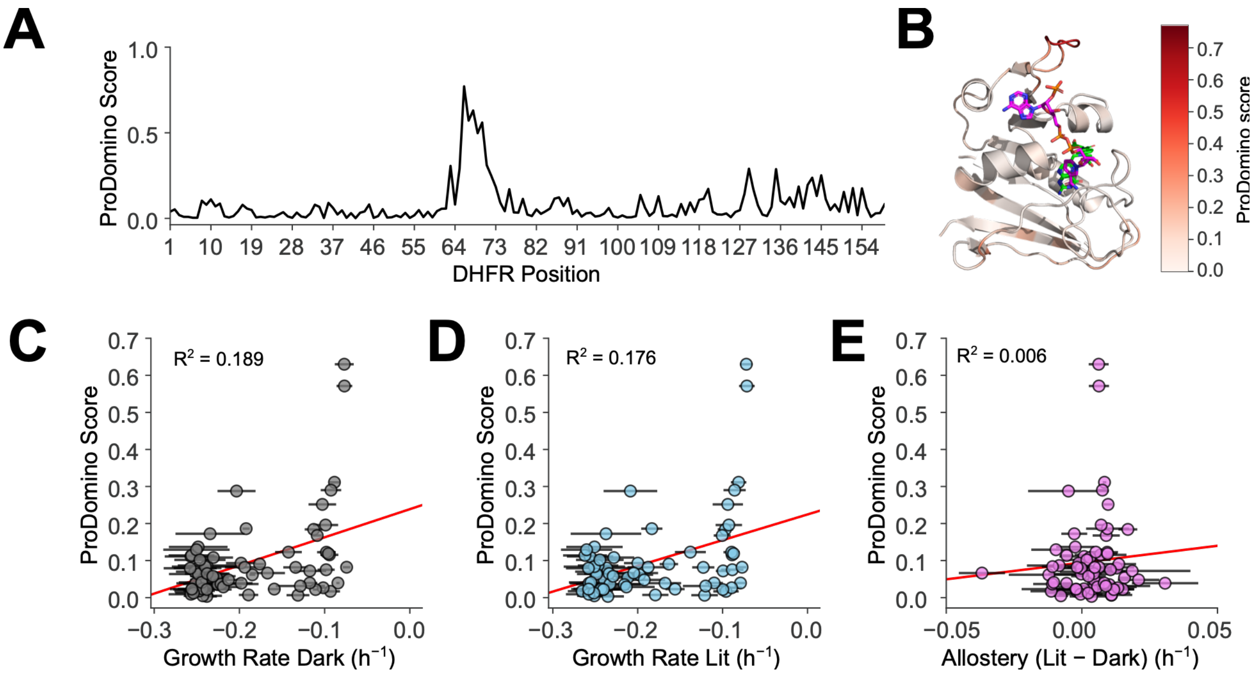


**Supplementary Figure S8: Comparison of DHFR-LOV2 insertion scan from Reynolds et al. 2011 to ProDomino predictions of domain insertion tolerance. (A)** ProDomino predictions of domain insertion tolerance for *E. coli* DHFR. **(B)** ProDomino scores visualized on the DHFR structure (PDB: 1RX2). **(C)** Scatterplot of ProDomino scores with growth rates of DHFR-LOV2 chimeras in the dark. **(D)** Scatterplot of ProDomino scores with growth rates of DHFR-LOV2 chimeras in the dark. **(E)** Scatterplot of ProDomino scores with allostery in DHFR-LOV2 chimeras. The red lines are lines of best fit.

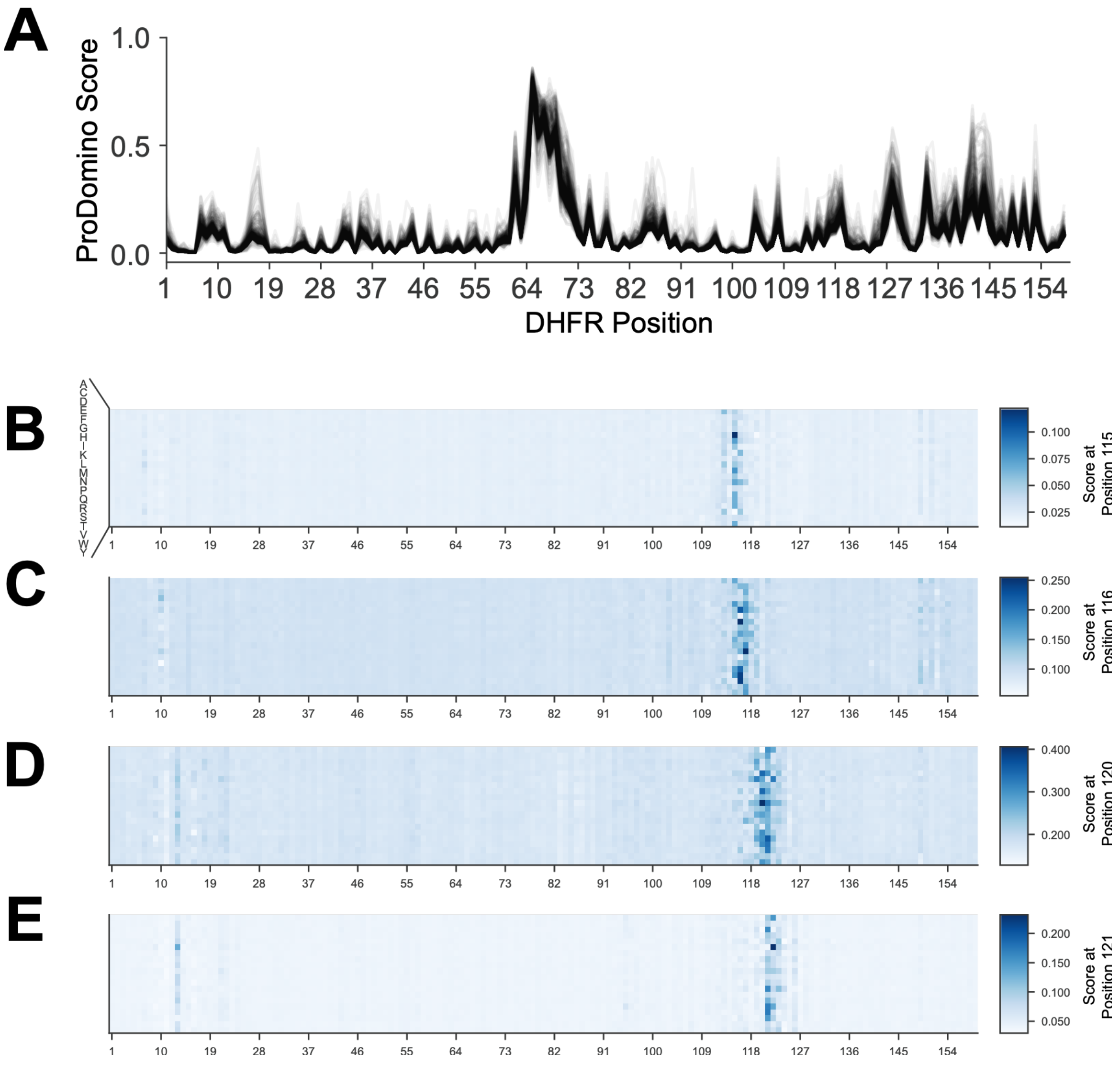


**Supplementary Figure S9: ProDomino predictions for DHFR single mutants. (A)** Overlaid ProDomino score profiles for single mutants of DHFR. **(B-E)** Heatmaps of ProDomino scores for sites (A) 115, (B) 116, (C) 120, and (D) 121 of DHFR in the background of each DHFR point mutant. These sites are the positions flanking the inserted LOV2 domain in DL121 and DL116. The vertical axes are the identity of the mutated residue, while the horizontal axes are the position in DHFR.

| | DL116 | | DL121 | |
|---|---|---|---|---|
| | **Dark** | **Lit** | **Dark** | **Lit** |
| **$k_{cat}$ (μmol $s^{-1}$)** | 0.38 ± 0.03 | 0.349 ± 0.001 | 0.33 ± 0.07 | 0.67 ± 0.07 |
| **$K_m$ (μM DHF)** | 1.5 ± 0.5 | 1.2 ± 0.1 | 1.5 ± 0.3 | 2.1 ± 0.3 |
| **m ($h^{-1}$)** | -0.34 ± 0.02 | -0.34 ± 0.01 | -0.32 ± 0.02 | -0.26 ± 0.01 |

**Table S1: Kinetic Parameters and growth rates of unmutated DL116 and DL121.** The values are shown as the mean plus or minus the standard deviation (n = 3 for kinetics, n = 4 for growth rates). The relative growth rates shown (m) are relative to WT DHFR and were re-fitted from McCormick et al. 2021 using weighted least squares linear regression.
